# Single-nucleus multiomics reveals the disrupted regulatory programs in three brain regions of sporadic early-onset Alzheimer’s disease

**DOI:** 10.1101/2024.06.25.600720

**Authors:** Andi Liu, Citu Citu, Nitesh Enduru, Xian Chen, Astrid M. Manuel, Tirthankar Sinha, Damian Gorski, Brisa S. Fernandes, Meifang Yu, Paul E. Schulz, Lukas M. Simon, Claudio Soto, Zhongming Zhao

## Abstract

Sporadic early-onset Alzheimer’s disease (sEOAD) represents a significant but less-studied subtype of Alzheimer’s disease (AD). Here, we generated a single-nucleus multiome atlas derived from the postmortem prefrontal cortex, entorhinal cortex, and hippocampus of nine individuals with or without sEOAD. Comprehensive analyses were conducted to delineate cell type-specific transcriptomic changes and linked candidate *cis-*regulatory elements (cCREs) across brain regions. We prioritized seven conservative transcription factors in glial cells in multiple brain regions, including RFX4 in astrocytes and IKZF1 in microglia, which are implicated in regulating sEOAD-associated genes. Moreover, we identified the top 25 altered intercellular signaling between glial cells and neurons, highlighting their regulatory potential on gene expression in receiver cells. We reported 38 cCREs linked to sEOAD-associated genes overlapped with late-onset AD risk loci, and sEOAD cCREs enriched in neuropsychiatric disorder risk loci. This atlas helps dissect transcriptional and chromatin dynamics in sEOAD, providing a key resource for AD research.

## Main

Sporadic early-onset Alzheimer’s disease (sEOAD), characterized by symptom onset before the age of 65 years^1^, represents a significant but under-studied type of Alzheimer’s disease (AD). While sEOAD shares neuropathological characteristics with late-onset AD (LOAD, symptom onset after the age of 65 years), including the extracellular accumulation of amyloid-β (Aβ) peptides and intracellular aggregation of tau proteins^2^, it often exhibits atypical presentations and accelerated disease progression^3^. Despite the phenotypic similarities, a substantial gap remains in the understanding of how alterations in gene expression and regulatory circuits in different brain regions contribute to the pathogenesis of sEOAD.

Recent advancements in single-nucleus RNA sequencing (snRNA-seq) and single-nucleus Assay for Transposase-Accessible Chromatin sequencing (snATAC-seq) have facilitated the examination of cell type-specific transcriptomic and epigenomic profiles in the human brain. These advancements have been paramount in elucidating the regulatory elements that govern cell type-specific gene expression patterns^4–6^. Furthermore, integrating snRNA-seq with snATAC-seq has proven invaluable in identifying candidate *cis-*regulatory elements (cCREs) and regulatory networks associated with LOAD^7–9^. However, comprehensive analyses of dysregulated regulatory circuits in the context of sEOAD remain limited. Importantly, previous research has primarily focused on regulatory mechanisms in the prefrontal cortex (PFC), but other two regions critically implicated in early stages of AD pathogenesis, the entorhinal cortex (EC) and hippocampus (HIP), have not been extensively studied^10, 11^.

Mutations in the *APP*, *PSEN1*, and *PSEN2* genes are known causes of Mendelian EOAD, but they account for only approximately 10% of EOAD cases^12, 13^. The remaining 90% of cases are classified as sEOAD, with their genetic underpinnings largely unidentified. Genetic studies in sEOAD have unveiled both common and rare variants in genes associated with LOAD, alongside novel variants, suggesting a partially overlapping genetic framework between sEOAD and LOAD^14–16^. Several genome-wide association studies (GWAS) have identified over 75 common variants linked to LOAD, with the *APOE* ε4 allele being the most significant genetic risk factor^17–22^. Moreover, studies have shown that most identified variants associated with LOAD reside in non-coding regions and are suggested to modulate gene expression through non-direct regulatory mechanisms in a cell type-specific manner^4, 7–9, 23^. Therefore, a systematic assessment of variants and their potential functions is required for understanding the genetic similarities and differences between sEOAD and LOAD.

To bridge these research gaps and delineate the molecular landscape of sEOAD, we generated single-nucleus multiome data of a small sEOAD cohort. The multiome profiling, which combines snRNA-seq and snATAC-seq in a single experiment, can effectively reduce batch effects among experiments. Our dataset includes gene expression and chromatin accessibility profiles from over 70,000 nuclei obtained from postmortem brain samples of the PFC, EC, and HIP regions of sEOAD individuals and age-matched control donors. Here, we conducted systematic bioinformatics analyses to detect cellular compositional changes, transcriptome alterations associated with sEOAD, and cell type-specific cCREs influencing gene expression across different brain regions. We examined transcription factors (TFs) that may exert their functions through distal enhancer-gene connections and identified their targeted genes in the brain of individuals with sEOAD. Furthermore, we examined the altered intercellular communications between non-neuronal and neuronal cell types and identified potential target genes in receiver cells in sEOAD. Finally, to explore shared and unique genetic components, we fine-mapped the genetic variants associated with LOAD and other traits to the cell type-specific chromatin accessible peaks in sEOAD brains. Altogether, our study provides a comprehensive overview of gene regulatory mechanisms in sEOAD brains, shedding light on the complex interplay between diverse brain cell types in its pathophysiology.

## Results

### Joint snRNA-seq and snATAC-seq profiling of sEOAD in three brain regions

To investigate the dysregulated gene regulatory mechanisms in sEOAD brains, we generated joint snRNA-seq and snATAC-seq (snMultiome) profiles from each sample. We conducted 10x Chromium single-nucleus multiome assays on the nuclei isolated from postmortem PFC (n = 9), EC (n = 6), and HIP (n = 5) tissues of the sEOAD individuals and matched controls. The four individuals with sEOAD (age at death: 59-64 years) did not have any established causal variants in *APP*, *PSEN1*, and *PSEN2* associated with EOAD, as documented on the Alzheimer’s Forum website (https://www.alzforum.org/mutations). The five age-matched controls (age at death: 61-65 years) had no neuropsychiatry diagnoses (**Fig. 1a & Supplementary Table 1**). Sample-level quality control was performed using the CellBender tool to remove systematic background noise in the snRNA-seq assay, followed by filtration of low-quality nuclei and putative doublets in both assays (**Supplementary Fig. 1, Methods**). In total, we generated high-quality transcriptomic and epigenomic profiles of 71,763 nuclei. The normalization, unsupervised dimension reduction, and integration were performed based on either gene expression, open chromatin accessibility profiles, or both modalities combined. The results revealed consistent cell clustering structures across three brain regions, as visualized using Uniform Manifold Approximation and Projection (UMAP) (**Fig. 1b, Supplementary Fig. 2**).

**Fig. 1:**
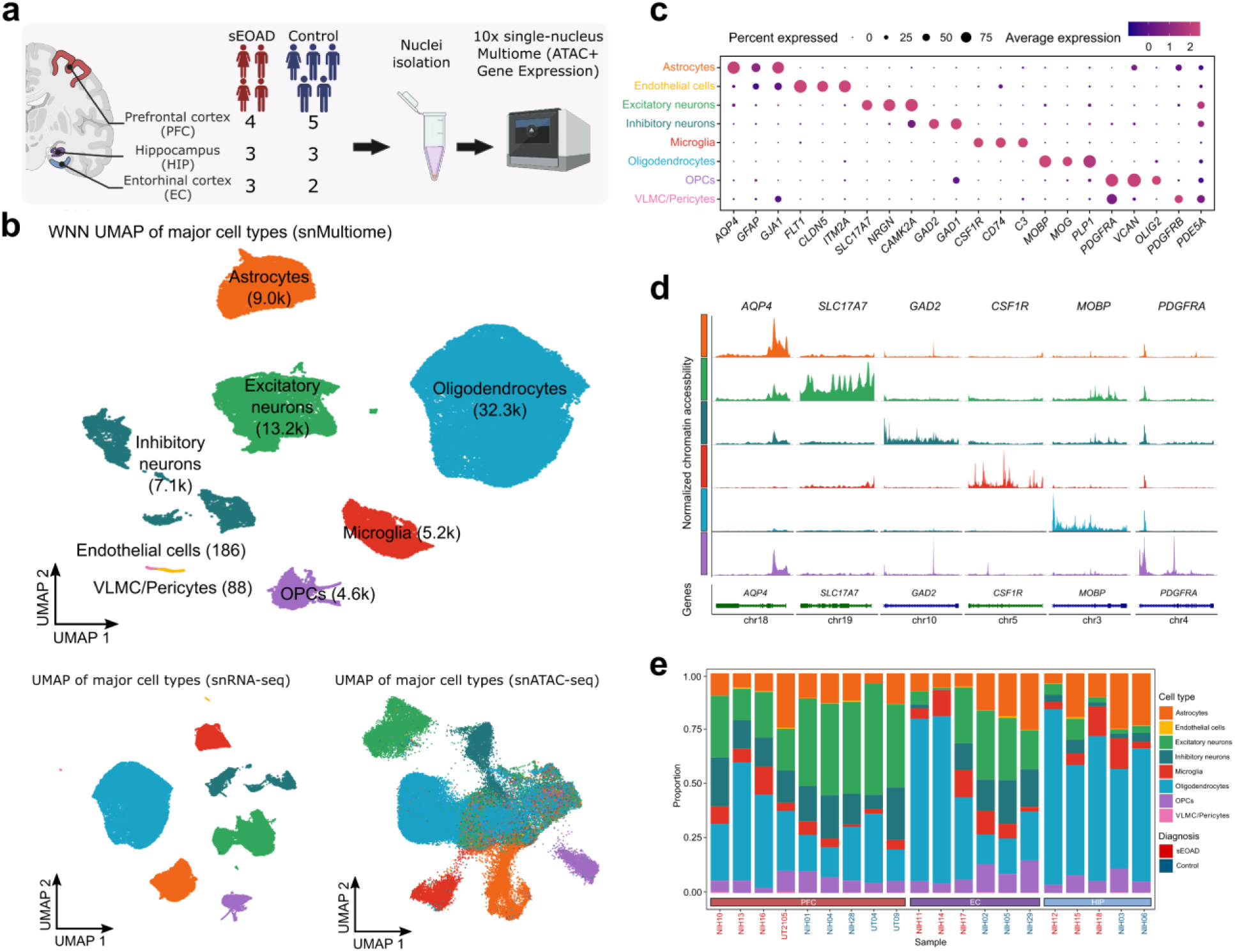
Single-nucleus multiome assays characterized cell type-specific expression and regulation in the three brain regions in sEOAD individuals. **a,** Experimental design (created through BioRender.com). **b,** Uniform manifold approximation and projection (UMAP) visualization of 71,763 nuclei based on weighted nearest neighbor (WNN) of both modalities (top), snRNA-seq assay (bottom left), and snATAC-seq assay (bottom right), colored by cell type. **c,** Dot plot illustrating selected marker gene expression (x-axis) across major cell types (y-axis). The color of the dots represents the scaled average expression of selected genes, and the size of the dots is proportional to the percentage of expressed cells. **d,** Chromatin accessibility of selected marker genes (x-axis) across major cell types (y-axis). Chromosome coordinates (human reference assembly GRCh38) are the following: *AQP4:* chr18:26851938-26866618; *SLC17A7:* chr19:49429301-49442860; *GAD2:* chr10:26215307-26304658; *CSF1R*: chr5:150053191-150113772; *MOBP*: chr3:39466198-39529579; and *PDGFRA*: chr4:54228097-54298347. **e,** The proportion of major cell types in each brain region of each sample. The color of the sample represents the diagnosis label.

We annotated the 71,763 nuclei by mapping the snRNA-seq profile to reference snRNA-seq datasets of the human brain from previous studies^24, 25^. These 71,763 nuclei were categorized into eight major cell types (with confidence score >0.95 from references) and high-resolution sub-cell types (**Fig. 1b, Supplementary Fig. 2a**). Consistent with previously published snRNA-seq studies of the human brain^7, 9, 24–26^, we observed cell type-specific marker gene expression within each annotated cluster, including *AQP4* in astrocytes, *SLC17A7* in excitatory neurons, *GAD2* in inhibitory neurons, *CSF1R* and *CD74* in microglia, *MOBP* in oligodendrocyte, *PDGFRA* in oligodendrocyte precursor cells (OPCs), *FLT1* in endothelial cells, and *PDE5A* in vascular and leptomeningeal cells (VLMC)/pericytes (**Fig. 1c**). Chromatin accessibility information around cell type marker genes also revealed unambiguous patterns distinguishing major cell types in the brain (**Fig. 1d, Supplementary Fig. 2b-d**). Together, these results indicated that the snRNA-seq and snATAC-seq data were of high quality and that cell type annotations were reliable. Because VLMC/pericytes and endothelial cells accounted for fewer than 200 nuclei, they were excluded from further analyses.

Our cell type composition analysis using the scCODA tool^27^ revealed the alterations in cell type abundance among different brain regions and conditions (**Fig. 1e, Extended Data Fig. 1, Supplementary Table 2**). Excitatory and inhibitory neurons were significantly less abundant in the EC and HIP compared to the PFC (posterior probability of inclusion >0.95), while the other six major cell types remained stable across regions (**Extended Data Fig. 1a**). These differences may be due to tissue sampling closer to the white matter in the HIP and EC, resulting in a higher proportion of glial cells compared to the PFC. Although a lower abundance of excitatory neurons was observed in the sEOAD PFC compared to controls, no significant differences in the proportion of other cell types were found between sEOAD and controls across the three brain regions (**Extended Data Fig. 1b-d**). These findings aligned with the notion that AD brains are pathologically characterized by neuronal loss, at least in the PFC^28^. Although not statistically significant, a higher abundance of oligodendrocytes was observed in sEOAD than in controls, particularly in the EC (**Extended Data Fig. 1b-d**).

### Cell type-specific dysregulation of gene expression in sEOAD across brain regions

To systematically characterize transcriptomic changes, we performed differential gene expression analysis in each major cell type of different brain regions between the sEOAD and control groups. We identified 1,222 sEOAD-associated differentially expressed genes (DEGs) in the PFC, 3,359 in the EC, and 5,430 in the HIP (**Supplementary Data 1**). These sEOAD DEGs were identified based on consensus signals from both MAST and mixed-effect models (adjusted p-value <0.05, absolute log2 fold change [|log2FC|] >0.25), accounting for sex, age at death, and batch (**Fig. 2a, Methods**). Despite similar numbers of nuclei captured in each brain region (**Supplementary Fig. 2b-d**), nearly three times as many sEOAD DEGs were found in the EC and HIP compared to the PFC. This finding from the snMultiome assay supports the previous reports that the EC and HIP are primarily affected in the early stages of AD pathogenesis^29^. Furthermore, sEOAD DEGs exhibited larger effect sizes (|log2FC| >1) in the HIP and EC compared to the PFC (**Fig. 2b**).

**Fig. 2:**
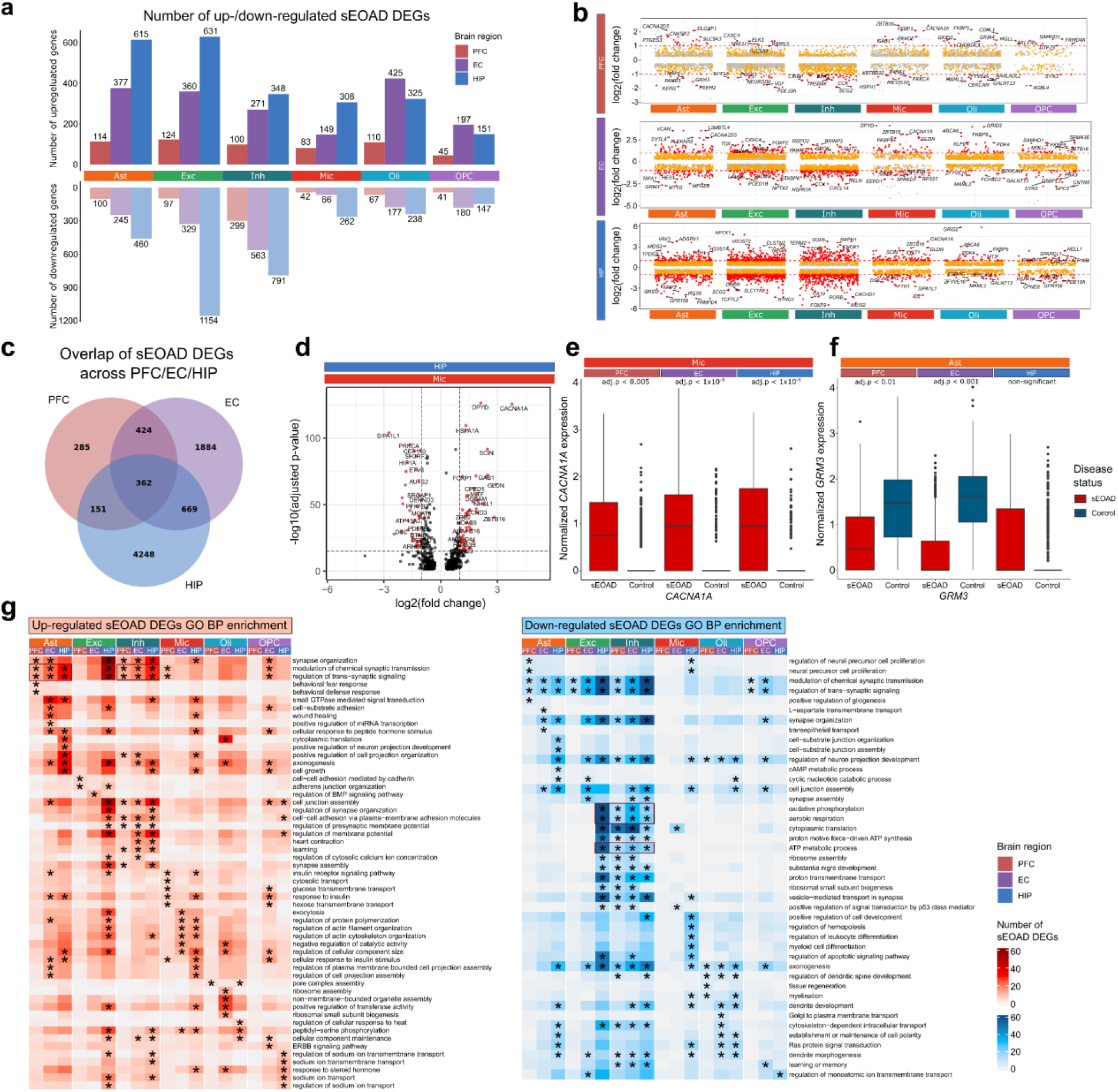
Cell type-specific transcriptomic alterations in sEOAD revealed dysregulation of synaptic signaling-related pathways in astrocytes and neurons across the brain regions. **a,** Number of up- and down-regulated differential expression genes (DEGs) in sEOAD (sEOAD DEGs) (y-axis) in each cell type in each brain region. **b,** Jitter plot highlighting the top sEOAD DEGs in each brain region with log2 fold change shown on the y-axis. The red dashed lines represent log2 fold change (log2FC) = 1. **c,** Venn diagram represents the number of the overlapped sEOAD DEGs in each cell type across brain regions. **d,** Volcano plot highlighting the top sEOAD DEGs of microglia in the hippocampus region. Gray vertical dashed lines represent the absolute log2FC = 1, and the gray horizontal dashed line represents the –log10 (adjusted p-value) = 16. **e,** *CACNA1A* gene was consistently upregulated in the microglia of sEOAD individuals across brain regions. **f,** *PTPRG* gene was significantly downregulated in astrocytes of the prefrontal cortex (PFC) and entorhinal cortex (EC) but upregulated in the hippocampus (HIP). The box boundaries and lines in the boxplot correspond to the interquartile range (IQR) and median, respectively. **g,** Heatmaps depicting the enriched Gene Ontology (GO) biological processes (BP) in up-(left) and down-(right) regulated sEOAD DEGs in different cell types of three brain regions.

Among the DEGs in the sEOAD PFC, 249 were consistent with those identified in previous snRNA-seq studies of LOAD PFC (Fisher’s exact test, p-value <2.2 ×10^−16^, **Extended Data Fig. 2a**). Notably, 184 DEGs were upregulated or downregulated in the same major cell types reported in the snMultiome study by Anderson et al.^9^. Three DEGs were consistent across all studies: upregulation of *MRAS* in astrocytes and upregulation of *CHORDC1* and *TP53TG5* in oligodendrocytes.

We identified 362 cell type-specific sEOAD DEGs consistently upregulated and downregulated across all three brain regions (**Fig 2c, Extended Data Fig. 2b**), including 88 genes previously reported in snRNA-seq studies of LOAD PFC^7, 9, 24^. For example, *CACNA1A* was consistently upregulated in microglia across the PFC, EC and HIP (**Fig. 2d, e, Supplementary Fig. 3**). *CACNA1A* encodes a subunit of a calcium-dependent voltage channel (CaV2.1). The upregulation of this gene, specifically in microglia, was reported to be associated with Aβ load in the human brain^30^. We also identified other consistently dysregulated genes, such as downregulated *CIRBP* and upregulated *ITGB8* in the astrocytes (**Extended Data Fig. 2c**). Those genes have been reported to be relevant to AD mechanisms^31, 32^.

Among the consistent DEGs across regions, we reported 274 genes that were not previously reported in snRNA-seq analysis of LOAD in the PFC. For example, *CXXC4*, which we found upregulated in excitatory neurons, is part of the Wnt signaling pathway and a key regulator of the amyloid precursor protein (**Extended Data Fig. 2c**). *MAML2*, which we found downregulated in oligodendrocytes, is part of the Notch signaling pathway, linked to neurovascular dysfunction in AD^33^. Additionally, 86 sEOAD DEGs were consistently downregulated in inhibitory neurons across three brain regions. Gene set enrichment analysis (GSEA) revealed their enriched Gene Ontology (GO) Biological Process (BP) terms, such as “modulation of chemical synaptic transmission”, “regulation of trans-synaptic signaling”, and “synapse organization” [Benjamini-Hochberg (BH) adjusted p-value <0.05, **Extended Data Fig. 2d**].

Most cell type-specific sEOAD DEGs showed varying dysregulation patterns across the considered regions. For example, *GRM3*, which encodes the glutamate metabotropic receptor 3 and is associated with synaptic function, was downregulated in astrocytes in the PFC and EC but not in HIP (**Fig. 2f**). This gene was found altered in AD and other neurological disorders^34, 35^.

In our GSEA of sEOAD DEGs for each cell type, we observed diverse enriched GO BP terms across the three regions (BH adjusted p-value <0.05) (**Fig. 2g**). Upregulated DEGs in astrocytes and inhibitory neurons were consistently enriched in synaptic functions, such as “modulation of chemical synaptic transmission” and “regulation of trans-synaptic signaling,” across all brain regions. However, only upregulated DEGs in excitatory neurons in the HIP were enriched in synaptic functions, indicating region-specific dysregulation. Upregulated genes in the microglia were enriched in cell morphology-related functions in the EC and HIP but not in the PFC. Downregulated DEGs in inhibitory neurons across all brain regions and in excitatory neurons in the HIP were enriched in “oxidative phosphorylation,” “aerobic respiration,” and “ATP metabolic process,” suggesting potential mitochondrial dysfunction and reduced energy metabolism in sEOAD brains^36, 37^.

### Cell type-specific candidate *cis-*regulatory elements (cCREs) for sEOAD DEGs

To assess potential cell type-specific open chromatin accessible signals associated with sEOAD, we performed peak calling on snATAC-seq assay using MACS2. We identified a total of 316,172 peaks in all major cell types, 75.3% of which were overlap with the results from a previously published cCRE dataset from the normal human brain by Li et al.^4^, and 61.5% of the peaks overlapped with the signals in the ENCODE cCREs database (**Fig. 3a**)^38^. Among these peaks, 22.45% were within 3 kbp of the nearest transcriptional start sites (TSSs), 46.21% were in intronic regions, and 24.13% were in distal intergenic regions (**Fig. 3b**).

**Fig. 3:**
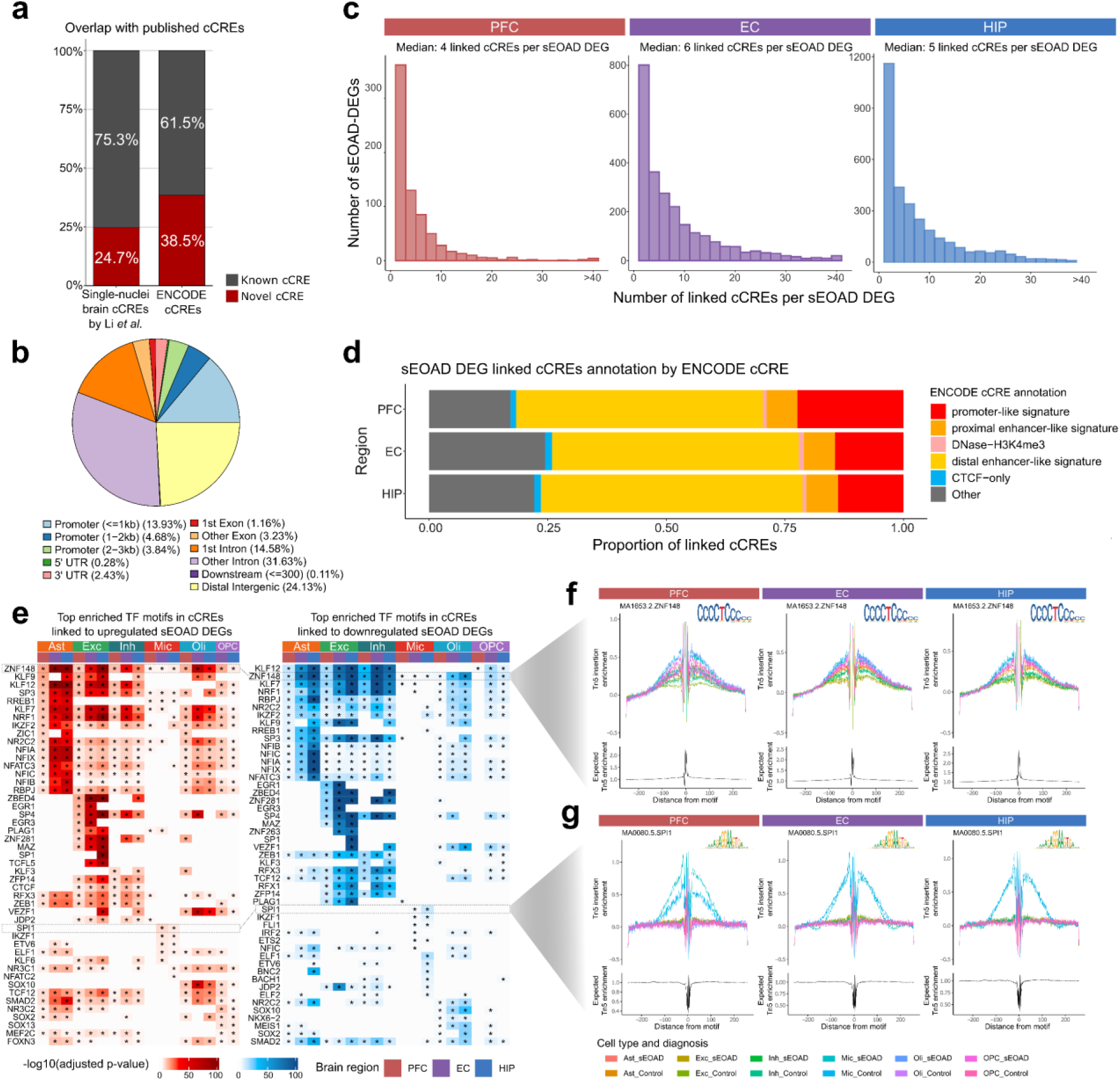
Identification of cis-regulatory elements linked to sEOAD DEGs in specific cell types and brain regions. **a,** Candidate *cis-*regulatory elements (cCREs) identified in each cell type. By comparing with the specific source, our cCREs were grouped into known and novel signals. **b,** Pie chart displaying the fraction of cCREs in genomic regions (anti-clock direction, starting with light blue for promoter). **c,** Density plots showing the distribution of the number of peaks linked to sEOAD differential expression genes (sEOAD DEGs) in each cell type in the prefrontal cortex (PFC, left), entorhinal cortex (EC, middle), and hippocampus (HIP, right), respectively. **d,** Stacked bar plot showing the percentage of peaks linked to sEOAD DEG in each brain region, overlapped with ENCODE cCRE annotated promoters and enhancers. **e,** Heatmaps depicting the enriched transcription factor (TF) motifs in cCREs linked to up-(left) and down-(right) regulated sEOAD DEGs in different cell types of three brain regions (* Benjamini-Hochberg adjusted p-value <0.05). **f,** and **g,** DNA footprinting analysis of candidate TFs: ZNF148 in all major cell types (**f**), and SPI1 in microglia (**g**). TF binding motifs are shown as a logo on the top-right corner of the panel.

To decode the dysregulated transcriptional regulatory circuitry in sEOAD brains, we conducted peak-to-gene linkage analyses in each brain region. We leveraged shared barcodes in snMultiome profiling to directly link epigenomic profiles with the respective transcriptomic profiles. We prioritized open chromatin peaks within 500 kbp of the TSS of each sEOAD DEG, focusing on significant positively correlated peaks (Spearman’s correlation >0.05, BH adjusted p-value <0.05) with the target gene, considering peak size, GC content, and fragment count. In total, we identified 3,794 cCRE-DEG associations in the PFC, involving 724 unique DEGs. In the EC, we found 21,349 of these associations, impacting 2,437 unique DEGs; while in the HIP, we reported 27,666 associations affecting 3,171 unique DEGs. Each cCRE was linked to a median of four DEGs in the PFC, six in the EC, and five in the HIP (**Fig. 3c**). The median distance between the cCRE and the TSS of the linked DEG was 140,980 bp in all three brain regions. The greatest proportion of the cCREs linked to sEOAD DEGs were present within distal enhancer-like regions based on the annotations of ENCODE cCRE database (52.00% in PFC, 51.99% in EC, and 55.11% in HIP) (**Fig. 3d, Supplementary Data 2**). This pattern was consistent in cCRE linked to sEOAD DEGs across cell types and brain regions (**Extended Data Fig. 3**). The results suggested the critical role of distal enhancers in gene expression alterations in sEOAD brains.

Transcription factors are the primary regulators of gene expression, and their dysregulation has been associated with AD pathogenesis^39^. Using snATAC-seq data, we conducted motif enrichment analysis to nominate TFs that may contribute to altered gene expression in sEOAD in each cell type of each brain region. To prioritize *bona fide* signals, we focused on TFs expressed in over 25% of the corresponding cell types. In total, 217 TF motifs were enriched in the cCREs linked to sEOAD DEGs (BH adjusted p-value <0.05, **Fig. 3e, Supplementary Data 3**). Among them, 148 motifs were significantly enriched in the cCREs linked to sEOAD DEGs across multiple cell types. For instance, motifs of ZNF148 had significant enrichment in cCREs associated with sEOAD DEGs in all major cell types (BH adjusted p-value <0.05, **Fig. 3e**). Other TFs, such as EGR1/3 in excitatory neurons and SPI1 and IKZF1 in microglia, appeared to exert their regulatory functions in specific cell types. The motif activity of these TFs within specific cell types and brain regions was further supported by DNA footprinting analyses (**Fig. 3f, g**). Importantly, most identified TF motifs (198 out of 217) were enriched in the cCREs positively correlated with the upregulated DEGs, and they also enriched in distinct cCREs correlated with downregulated DEGs in the same cell type. This suggested TFs may function as both activators and repressors in different chromatin regions, thereby targeting different genes. Indeed, it has been reported that the effect of TFs on transcription is context-dependent; TFs can recruit different coactivators or corepressors, forming multi-subunit protein complexes to regulate transcription^40, 41^.

### Cell type-specific regulatory networks in the sEOAD brain

To further explore the alterations in sEOAD brains, we identified cell type-specific TFs potentially modulating gene expression through enhancer binding in each brain region (**Fig. 4a**). Using SCENIC+ algorithm, we inferred enhancer-driven regulons for each brain region, which comprising candidate TFs, open chromatin regions with enriched TF-binding motifs, and target genes across cellular states^42^. We prioritized the regulons exhibiting the highest 5% positive correlations and the lowest 5% negative correlations between TF expression and open chromatin region-based regulon enrichment scores. We identified 16, 31, and 17 key regulators in the PFC, EC, and HIP, respectively (**Fig. 4b, Extended Data Fig. 4a, b, Supplementary Data 4**). These regulators may act as either activators or repressors. On average, each regulon comprised 1,014 open chromatin regions and 262 target genes. We observed high cooperativity in prioritized regulons, as evidenced by the shared target regions and genes among them (**Extended Data Fig. 4c-e**). For instance, the chromatin regions and genes targeted by IKZF1 and FLI1 were highly overlapped in all three brain regions (Jaccard coefficient >0.5, **Extended Data Fig. 4c-e**). These results suggested that a complex regulatory network, involving cooperativity from multiple TFs, is necessary to influence the transcriptome profiles in the human brain^43, 44^. Seven regulators were found to be conserved in specific cell types across all three brain regions, including RFX4 in astrocytes and FLI1, IKZF1, RUNX1, FOXN3, ETS2, and POU2F2 in microglia (**Extended Data Fig. 4f**).

**Fig. 4:**
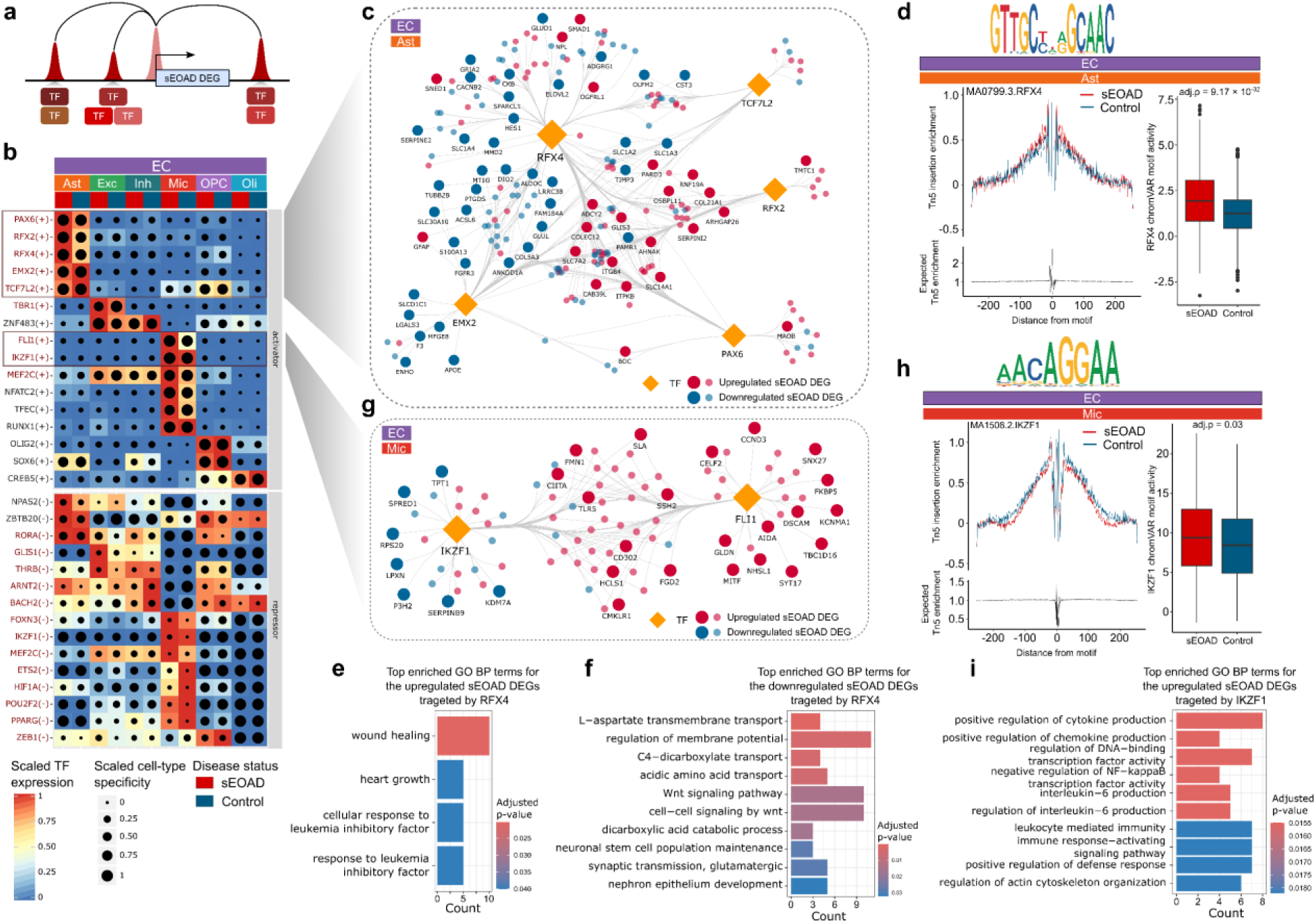
Cell type-specific, dysregulated transcription factor (TF) regulatory circuities in the EC in sEOAD. **a,** Illustration of TF binding to a target gene. **b,** Heatmap/dot plot depicting key regulon of the entorhinal cortex (EC) by cell type and diagnosis label (x-axis). The expression of each TF is presented on a color scale, and the cell type-specificity of the regulon is shown on a dot-size scale. **c,** The reconstructed TF regulatory network indicating RFX4, PAX6, RFX2, EMX2, and TCF7L2 regulate sEOAD DEGs in astrocytes within the EC region. **d,** Left: DNA footprinting analysis for RXF4 in astrocyte within EC by sEOAD diagnosis label. Right: Box plot of chromVAR motif activity of RFX4 shows a significant difference in astrocyte between sEOAD and control. TF binding motif is shown as a motif logo on the top of the panel. Box boundaries and lines correspond to the interquartile range (IQR) and median, respectively. **e** and **f,** Top enriched Gene Ontology (GO) Biological Process (BP) terms for up-(**e**) or down-(**f**) regulated sEOAD DEGs targeted by RFX4 in astrocytes within EC. **g,** The reconstructed TF regulatory network showing IKZF1 and FLI1 regulate sEOAD DEGs in microglia within the EC region. **h,** Left: DNA footprinting analysis for IKZF1 in microglia within EC by sEOAD diagnosis label. Right: The box plot of chromVAR motif activity of IKZF1 shows a significant difference in microglia between sEOAD and control groups. TF binding motif is depicted as a motif logo on the top of the panel. **i,** Top enriched GO BP terms in upregulated sEOAD DEGs targeted by IKZF1 in microglia of the EC region.

We further explored the gene regulatory networks in the EC region. SCENIC+ identified 31 regulons, indicating high regulatory activity (**Fig. 4b**). Among these regulons, we prioritized TFs with the motifs significantly enriched in cCREs linked to sEOAD DEGs in the corresponding cell types (**Fig. 3e, Supplementary Data 3**). This process pinpointed five activators in astrocytes (RFX4, PAX6, RFX2, EMX2, and TCF7L2) and two in microglia for constructing gene regulatory networks. The five activators in astrocytes targeted a total of 102 upregulated and 113 downregulated DEGs in the EC (**Fig. 4c, Extended Data Fig. 4e**). Particularly, RFX4 showed high specificity and regulatory activity in astrocytes across sEOAD and control samples, as validated by DNA footprinting analysis (**Fig. 4d**). The inferred chromVAR motif activity of RFX4 was significantly higher in astrocytes of sEOAD individuals (BH adjusted p-values = 9.17 ×10^−32^, **Fig. 4d**), indicating altered regulatory activity^45^. The RFX4 regulon contained 661 target genes, including 82 genes upregulated and 96 genes downregulated in astrocytes of sEOAD individuals. RFX4 (Regulatory Factor X 4) may play a role in psychiatric conditions in regulating genes crucial for brain development and function, including those involved in circadian rhythms and neural differentiation^46^. However, the disrupted regulatory effect of RFX4 on the transcriptome alterations in astrocytes of sEOAD has not been fully explored. In our sEOAD data, we observed the upregulation of *GFAP,* which encodes for glial fibrillary acidic protein, which is associated with astrocyte activation during stress and injury^47^. The GSEA of the DEGs targeted by RFX4 in astrocytes revealed diverse enriched GO BP terms, including “L-aspartate transmembrane transport”, “regulation of membrane potential”, and “Wnt signaling pathway”, among others (BH adjusted p-values <0.05, **Fig. 4e, f**). This signified its potential role in regulating multiple transport-related pathways in the brain of sEOAD.

Furthermore, we prioritized activators in the microglia of EC, including FLI1 and IKZF1. Both TFs exhibited high specificity within microglial cells, as their motifs were significantly enriched in cCREs linked to sEOAD DEGs in microglia. Specifically, these TFs targeted 71 upregulated and 20 downregulated DEGs (**Fig. 4g**). DNA footprinting analysis of IKZF1 showed high motif activity in the microglia of both sEOAD and control samples. Moreover, the inferred chromVAR motif activity was notably increased in sEOAD (adjusted p-value =0.03, **Fig. 4h**). The IKZF1 regulon involved 47 upregulated and 20 downregulated DEGs in the microglia of sEOAD. The dysregulated genes targeted by IKZF1 were predominantly involved in immune-related functions such as “negative regulation of immune response”, “leukocyte-mediated immunity”, and “immune response-activating signaling pathways” (BH adjusted p-value <0.05, **Fig. 4i**). This underscores the critical role of IKZF1 in mediating the immune response and pathophysiology of sEOAD.

Collectively, these results suggested that the cell type-specific transcriptomic profiles were regulated through cooperative interactions of multiple TFs and enhancers on their target genes in sEOAD brains. We highlighted a disrupted regulatory role for RFX4 in the astrocytes of sEOAD EC, which modulates genes involved in intercellular signaling pathways, and for IKZF1 in microglia, related to innate immunity and neuroinflammation.

### Dysregulated intercellular signaling between non-neuronal and neuronal cell types

The altered regulatory relationships in non-neuronal cell types and their impact on neuronal circuit homeostasis and pro-inflammatory responses might play an important role in LOAD brains ^48^. To estimate such association in sEOAD brains, we conducted comparative cell-cell communication analyses using the MultiNicheNet tool^49^. This analysis facilitated the identification of dysregulated intercellular communication mediated by ligand-receptor pairs and their potential regulatory influence on the receiver cells. We prioritized the top 25 differential signals sent and received by specific cell types as condition-specific intercellular signaling.

Here, we highlighted the intercellular communication analysis of astrocytes and microglia in the EC. Specifically, the results showed that all top 25 differential signals sent by astrocytes were downregulated in sEOAD (**Fig. 5a**). The communication signals between EFNA5 in astrocytes to EPHA3 and EPHA6 in excitatory neurons were downregulated (**Fig. 5a**). The Ephrin–Eph signaling plays an important role in regulating cell migration in neurons. It is also associated with cytoskeletal rearrangements, as it regulates small GTPase activation and inactivation^50^. Furthermore, we observed multiple downregulated ligand-receptor pairs involved in neuroinflammation regulation, such as PTN-PTPRS/PTPRB signaling from astrocytes to excitatory neurons and A2M-LRP1 signaling from microglia to astrocytes (**Fig. 5a, b**). Remarkably, although more upregulated intercellular signalings were observed in the sEOAD PFC, downregulated PSAP-LRP1 signaling from microglia to astrocytes was noted in all three brain regions (**Extended Data Fig. 5a-d**). The reduced levels of PSAP protein have been associated with increased neurofibrillary tangle development and neuroinflammation^51^. Furthermore, we identified upregulated TGFB1-ITGB8 signaling from microglia to astrocytes, suggesting a potential increasing microgliogenesis in the sEOAD EC and PFC^52^.

**Fig. 5:**
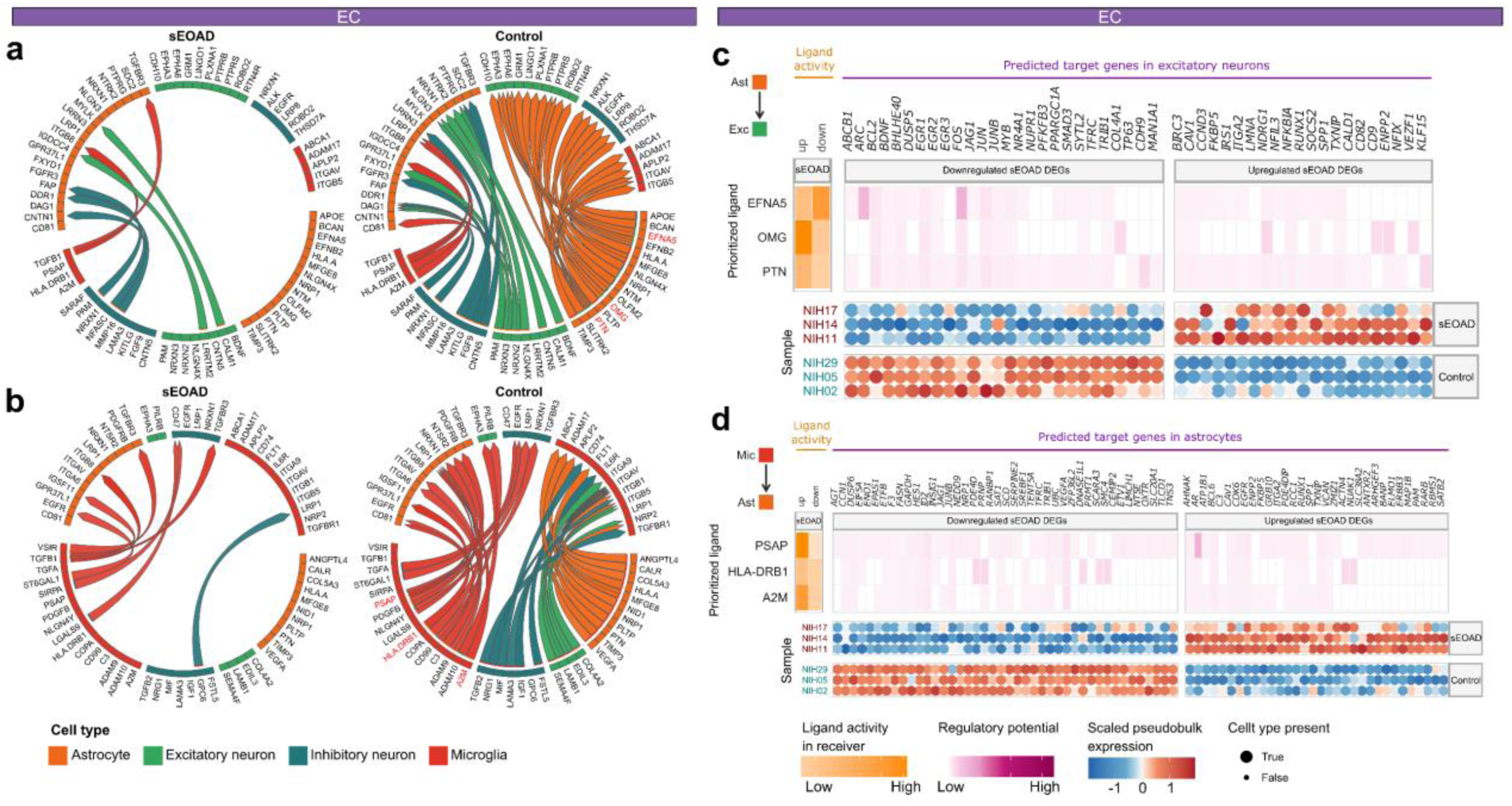
Comparative intercellular signaling analysis reveals dysregulated cell-cell communication signals between sEOAD and controls in the EC. **a,** Top 25 differential ligand-receptor signals received and sent by astrocytes in the entorhinal cortex (EC) between the sEOAD and control groups. **b,** Top 25 differential ligand-receptor signals received and sent by microglia in the EC between the sEOAD and control groups. The arrowhead indicates the direction of the signal from the sender cell type to the receiver cell type. **c,** Heatmap depicting the scaled (purple) and raw (orange) regulatory potential scores of selected ligands from dysregulated ligand-receptor pairs in astrocytes to differentially expressed genes (DEGs) in excitatory neurons. The dot plots represent the scaled pseudobulk expression of DEGs targeted by selected ligands, with dot size indicating whether a sample contained at least ten cells for excitatory neurons. **d,** Heatmap depicting the scaled (purple) and raw (orange) regulatory potential scores of selected ligands of dysregulated ligand-receptor pairs in microglia to sEOAD DEGs in astrocytes. The dot plots show the scaled pseudobulk expression of sEOAD DEGs targeted by selected ligands, with dot size indicating whether a sample contained at least ten cells for astrocytes.

We further explored the regulatory influence of ligands in sender cell types on the sEOAD DEGs in receiver cell types. Among the top differential communication pathways from astrocytes to excitatory neurons, ligands encoded by the *EFNA5*, *OMG*, and *PTN* in astrocytes exhibited increased ligand activity (**Fig. 5c**). Ligand EFNA5 exhibited the highest regulatory potential on downregulated expression of *ARC*. The Activity-Regulated Cytoskeleton-associated protein encoded by *ARC* plays a significant role in synaptic plasticity and neuronal activity^53^. Additionally, the TF coding genes in excitatory neurons, such as *EGR1*, *EGR3* and *FOS*, were targeted by ligands in astrocytes (**Fig. 5c**). Similarly, the ligands EFNA5 and PTN in astrocytes displayed elevated regulatory activities on downstream genes in excitatory neurons in the PFC (**Extended Data Fig. 5e**). Conversely, ligands encoded by *PSAP* and *A2M* in microglia exhibited high ligand activity, potentially modulating the expression of targeted genes in astrocytes (**Fig. 5d, Extended Data Fig. 5e, f**). These results underscored the potential subtle and dynamic intercellular signaling influences on neuronal and glial functions in sEOAD brains.

### Genetic variation at candidate CREs linked to sEOAD DEGs

To better understand the cellular impact of genetic variants identified by AD GWAS, we prioritized risk loci that overlapped with cCREs linked to sEOAD DEGs. We curated 148 index single nucleotide polymorphisms (SNPs), reported as genome-wide significant from four LOAD GWA studies (**Fig. 6a**)^17, 18, 20, 21^. We expanded these SNPs using linkage disequilibrium (LD) to include nearby variants with high coinheritance probability (LD R^2^ >0.8 based on phase 3 of the 1000 Genomes Project data). We identified 3,180 unique SNPs associated with LOAD, of which 98.2% were in non-coding regions. Interestingly, 133 LOAD SNPs mapped in cCRE sequences were linked to 29 sEOAD DEGs across each brain region (**Extended Data Fig. 6a**). Particularly, cCREs in microglia in the EC and HIP were significantly enriched with LOAD GWAS variants, unlike those in the PFC (adjusted Fisher’s exact test, p-value <0.05, **Fig. 6b**). In addition, cCREs linked to sEOAD DEGs of excitatory neurons in HIP were enriched with LOAD GWAS loci. Notably, cCREs linked to the downregulated *HLA-DRB1* and *HLA-DRA* genes in microglia intersected with the largest number of SNPs associated with LOAD (**Extended Data Fig. 6a**). The index SNP rs9271058 (reported in Kunkle et al.^18^) was mapped to a cCRE (chr6:32607513-32607728) linked to *HLA-DRB1* in microglia in the EC (**Fig. 6c**). There were 43 SNPs mapped to a cCRE (chr6:32603881-32604799) that was linked to *HLA-DRB1*. Importantly, this distal-like enhancer cCRE contains TF motifs specific to microglia regulators, such as IKZF1 and FLI1. We found that the index SNP (rs74685827), and its associated SNPs reported by Bellenguez *et al*.^21^, intersected with the microglia- and astrocytes-specific cCREs linked to dysregulated *SORL1* gene expression in the HIP **(Extended Data Fig. 6b**). We extended this approach to include 172 unique SNPs (LD R^2^ >0.8) from five index SNPs associated with EOAD (p-value <5×10^−8^)^54^. We pinpointed 20 SNPs that intersected with cCREs correlated with *HLA-DRB1* and *MS4A6A* in microglia within the EC, and 16 SNPs intersected with cCREs linked to *HLA-DRB1* and *HLA-DRA* in the HIP. We further examined whether the cCREs of sEOAD DEGs harbored single-cell expression quantitative trait locus (sc-eQTL) identified in the human brain^55^. We identified 6, 27, and 33 brain sc-eQTL loci intersecting with cCREs of sEOAD DEGs in the PFC, EC, and HIP, respectively (**Extended Data Fig. 6c, d**).

**Fig. 6:**
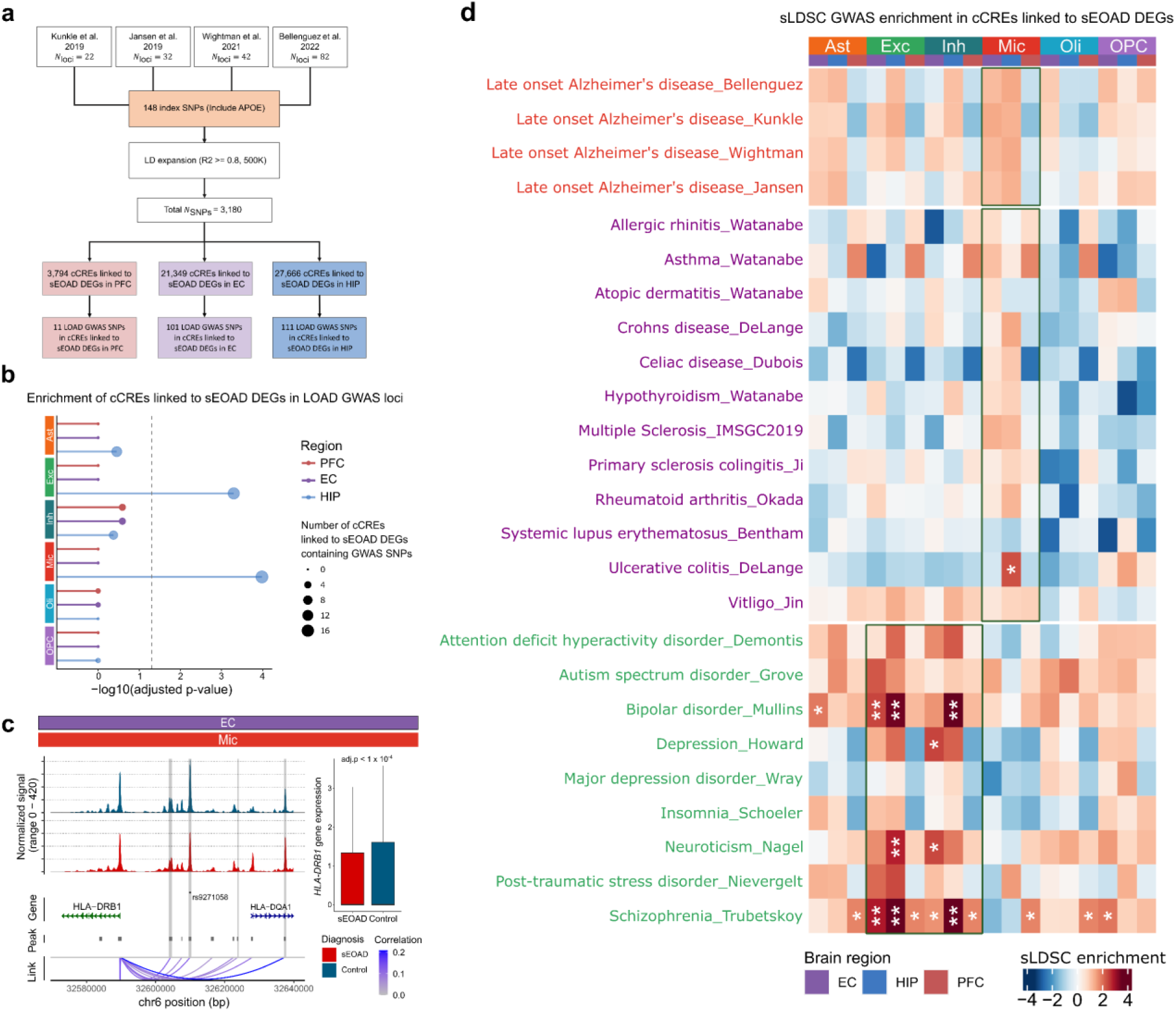
Mapping genome-wide associated loci with cell type- and region-specific cCREs in sEOAD. **a,** Candidate *cis-*regulatory elements (cCREs) linked to sEOAD differential expression genes (sEOAD DEGs) overlapped with single nucleotide polymorphisms (SNPs) associated with late-onset Alzheimer’s disease (LOAD). **b,** cCREs linked to sEOAD DEGs in microglia in the entorhinal cortex (EC) and hippocampus (HIP) were significantly enriched with LOAD GWAS identified SNPs. The p-values were calculated based on Fisher’s exact test. The dot size indicates the number of cCREs linked to sEOAD DEGs that harbor LOAD GWAS loci. **c,** The cCREs linked to *HLA-DRB1* overlapped with SNPs identified in LOAD GWAS, including the index SNP rs921058. The box plot shows that the *HLA-DRB1* gene was significantly down-regulated in microglia in the EC region of sEOAD. The cCREs linked to sEOAD DEGs harbor LOAD GWAS loci are highlighted in gray. **d,** Heatmap depicting the enrichment of risk variants associated with LOAD, 12 immune-mediated traits, and 9 psychiatry disorders from GWAS in cCREs linked to sEOAD DEGs in each cell type and brain region (*FDR <0.05, **FDR <0.01).

We next performed stratified-linkage disequilibrium score regression (sLDSC)^56^ to assess the heritability enrichment of genetic risk loci in cCREs linked to sEOAD DEGs in each cell type. In this analysis, we considered risk loci identified by LOAD, immune-mediated disorders, and other psychiatric disorders GWAS ^57–74^ (**Supplementary Tables 3**). Strikingly, we did not find enrichment of the LOAD-related SNPs in cCREs linked to sEOAD DEGs (BH adjusted p-value >0.05, **Fig. 6d, Supplementary Data 5a**). However, cCREs linked to sEOAD DEGs in microglia in the HIP were significantly enriched with risk loci of ulcerative colitis (BH adjusted p-value <0.05), though other immune-mediated traits did not show significant enrichment after multiple testing corrections. Conversely, we found significant heritable enrichment for neuropsychiatric disorders in cCREs linked to sEOAD DEGs across brain regions. Bipolar disorder and schizophrenia risk variants were significantly enriched within cCREs linked to sEOAD DEGs of excitatory and inhibitory neurons (BH adjusted p-value <0.05, **Fig. 6d**).

Similarly, we performed sLDSC analysis on cell type-specific cCREs (**Methods**). As expected, microglia-specific cCREs demonstrated significant enrichment for the heritability of LOAD, as well as various immune-mediated disorders, such as Crohn’s disease, multiple sclerosis, rheumatoid arthritis, ulcerative colitis, and vitiligo (adjusted p-value <0.05, **Extended Data** Fig. 6e**, Supplementary Data 5b**). A recent study revealed shared and unique genetic components among these disorders^75^. Furthermore, cell type-specific cCREs in excitatory neurons were significantly enriched in risk variants associated with neuropsychiatric disorders.

In summary, the cell type-specific cCREs linked to sEOAD DEGs suggested that LOAD risk variants in open chromatin peaks may account for a limited number of microglia transcriptome alterations. Further research is needed to determine the distinct genetic risk factors related to sEOAD.

## Discussion

We reported the first single-nucleus multiome profiling of sEOAD, creating a comprehensive atlas of gene expression and regulation in the PFC, EC, and HIP regions. By profiling 71,163 nuclei from four sEOAD cases and five matched controls, we simultaneously assessed chromatin accessibility and gene expression. Our analyses identified cell type-specific transcriptome alterations in sEOAD and linked cCREs across brain regions. We characterized conserved regulons in non-neuronal cells and found that multiple TFs may cooperatively modulate transcriptomic changes, leading to dysregulated synaptic plasticity and neuroinflammation in astrocytes and microglia. Additionally, we prioritized altered intercellular signaling pathways and their regulatory impacts on sEOAD DEGs. Our genetic fine-mapping analysis revealed cCREs linked to sEOAD-associated genes intersected with late-onset AD risk loci and may share genetic risks with neuropsychiatric disorders.

The inclusion of multiple brain regions in our study has enabled us to examine conserved regulators and their target genes across the brain regions that are disrupted in sEOAD. Our results highlighted regulators with high specificity in glial cells, such as RFX4 in astrocytes and IKZF1 in microglia. Remarkably, the RFX4 motif was enriched in cCREs linked to sEOAD DEGs across all regions, with targeted DEGs consistently enriched in transmembrane transport and trans-synaptic signaling functions. This suggests a conserved role for RFX4 in astrocytes and its universal regulatory alterations in sEOAD pathogenesis. Similarly, in microglia, upregulated genes related to neuroinflammation were targeted by TFs like IKZF1 and FLI1 in sEOAD brains. Additionally, we found cCREs contain TF motifs specific to microglia regulators, such as IKZF1 and FLI1, suggesting that SNPs in these cCREs may impact gene expression regulation to confer sEOAD risk. Our findings indicate that gene expression changes in sEOAD are regulated by complex synergistic and/or antagonistic effects of multiple TFs through distal enhancers.

We further prioritized cCREs linked to sEOAD DEGs in each brain region through integrated epigenomic and transcriptomic analysis. Interestingly, we found that cCREs linked to sEOAD DEGs in microglia intersected with genetic risk loci identified in LOAD GWAS studies^17, 18, 20, 21^. However, we did not find significant heritability enrichment in the sLDSC analysis (**Fig. 6d**). Genetic variants associated with psychiatric disorders and traits, such as schizophrenia, bipolar disorder, and neuroticism, were significantly enriched in both the cCREs linked to sEOAD DEGs and cell type-specific cCREs in excitatory neurons. This suggested potential shared genetic risks between sEOAD and psychiatric conditions, particularly in excitatory and inhibitory neurons^4, 7^. Our integrated epigenomic and transcriptomic analysis successfully explored the shared and unique genetic components and their potential functions associated with sEOAD.

Our study offers insights into cell type- and brain region-specific transcriptome dysregulation and potential regulatory disturbances in sEOAD. However, our study has limitations. First, this study is constrained by a small sample size, resulting from the limited availability of postmortem brain samples in sEOAD. We conducted a simulation-based power analysis and found lower statistical power estimated for identifying differentially expressed genes in less abundant cell types, such as inhibitory neurons (**Supplementary Table 4**). This limitation, due to the availability of brain tissue and also the high cost in multiome, also restricts our ability to perform further analyses at the sub-cell type level. We anticipate that larger-scale single nuclei studies will enhance the accuracy and detail in identifying gene expression changes in sEOAD. Second, future laboratory experiments should help validate the candidate TFs and cCREs in specific cell types. Third, our study did not comprehensively assess the regulatory mechanisms involving the complex interactions of multiple regulators on target genes. Finally, we lacked the access to sEOAD GWAS datasets for combined association with our scMultiome features, since they are not publicly available yet. We will integrate such analysis when sEOAD GWAS datasets are released to the public.

In conclusion, our study provides novel insights into transcriptome alterations and identifies key gene regulators, such as RFX4 in astrocytes and IKZF1 in microglia, that are crucial in the pathophysiology of sEOAD across three brain regions commonly affected in sEOAD. The intercellular communication analysis revealed the altered roles of non-neuronal cell types in synaptic plasticity, neuroinflammation, and microgliogenesis in sEOAD. We delineated distinct genetic architectures within cell type-specific cCREs. These discoveries set the stage for future investigations into the unique mechanisms of sEOAD pathophysiology and for the development of targeted therapeutic strategies.

## Methods

### Human postmortem brain sample collection

Postmortem brain samples of nine individuals were obtained from the NIH NeuroBioBank from the University of Miami and McGovern Medical School at UTHealth (**Supplementary Table 1**) in compliance with UTHealth Houston’s Institutional Review Board. sEOAD cases from the McGovern Medical School were defined as subjects diagnosed with sEOAD before the age of 65 years and confirmed neuropathological diagnosis postmortem. Brain samples of sEOAD from the NIH NeuroBioBank were selected based on the neuropathological diagnosis of sEOAD with onset before age 65 years. The genotype of the samples from the NIH NeuroBioBank were examined based on paired whole-genome sequencing data publicly available on The National Institute of Mental Health Data Archive (https://nda.nih.gov/edit_collection.html?id=3917). Subjects with known autosomal dominant mutations in *APP*, *PSEN1*, and *PSEN2* were excluded from the analyses. The sEOAD cases from McGovern Medical School were the subjects diagnosed with sEOAD before the age of 65 years and confirmed neuropathological diagnosis postmortem. All tissues were de-identified under the privacy rules of the Health Insurance Portability and Accountability Act of 1996 (HIPAA). The tissue donation consent was obtained from all participants by UTHealth Institutional Review Board.

### Tissue preparation and nuclei isolation

The tissue preparation, dissociation, and nuclei extraction were performed at the UTHealth Cancer Genomic Core. Samples were stored at −80 °C. The nuclei isolation protocol was modified based on a published protocol^76^ and 10x Genomics protocol (CG000375). Briefly, 30 mg of frozen tissue was finely minced on dry ice and then homogenized in buffer on ice within 20 minutes. The homogenized tissue was filtered through a 35 μm cell strainer. Small cell debris was removed by centrifuging at 300*g* for 3 minutes at 4 °C. The large cell debris was removed by gradient centrifugation using OptiPrep density gradient medium (Sigma, cat# D1556-250ML) at 7200*g* for 15 minutes at 4 °C. The nuclei pellet was resuspended in a 0.1x lysis buffer. After washing the isolated nuclei in the wash buffer, the nuclei were resuspended in 1x nuclei buffer before single-nuclei Multiome library construction.

### Single-nuclei multiome sequencing

Single-nuclei libraries were constructed following the 10x Chromium Single Cell Multiome ATAC + Gene Expression protocol (CG000338). Briefly, nuclei suspensions were incubated with a transposase, which fragmented the DNA in open regions of the chromatin and added the adapter sequences to the ends of the DNA fragments. The transposed nuclei were loaded onto Chromium Next GEM Chip J (PN-1000234, 10x Genomics, Pleasanton, CA) with portioning oil and barcoded single-cell gel beads, followed by PCR amplification. The ATAC and the gene expression libraries were then prepared separately. Library qualities were assessed using the Agilent High Sensitive DNA Kit (#5067-4626) on an Agilent Bioanalyzer 2100 (Agilent Technologies, Santa Clara, USA). Qualified libraries underwent paired-end sequencing on an Illumina NovaSeq System (Illumina, Inc., USA).

### Joint single-nuclei multiome data processing and cell type annotation

We utilized the Cell Ranger ARC (v2.0.2) pipeline for aligning the snRNA-seq and snATAC-seq reads from each sample to the GRCh38 genome and quantifying them with ‘*cellranger-arc count*’. The raw snRNA count matrix and snATAC fragment counts were further processed using a rigorous pipeline. Firstly, we removed the systematic background noise in the raw snRNA count matrix using CellBender, which applies a deep-learning model to accurately distinguish cell-containing droplets from cell-free droplets^77^. Secondly, we applied initial filtration based on RNA-assay matrices (200 < nFeature_RNA < 7,500) and ATAC-assay matrices (Nucleosome_signal < 2.5, TSS.enrichment > 2) using Seurat (v5.0.0)^78^ and Signac (v1.12.0)^79^ packages, respectively. Potential doublets were removed using the DoubletFinder R package using default parameters^80^. Then, the RNA counts were normalized using SCTransform with mitochondrial percent per cell regressed out. Principal component (PC) analysis (PCA) and UMAP visualization based on the first 30 PCs were performed on normalized RNA counts. Additionally, we performed pseudo-bulk analysis on RNA assay to identify and remove outlier samples through PCA.

Predicted major cell type annotations were determined for each cell based on SCTransform normalized reference mapping embedded in Seurat. Gene expression (RNA assay) was mapped to SCTransformed normalized snRNA-seq datasets of the PFC and middle temporal gyrus^24, 25^ and annotated with the cell types of the reference map. Cells with consistent cell type predictions and predicted cell type scores greater than 0.95 in both datasets were kept for downstream analysis. Additionally, we leveraged scANVI from scvi-tools (v1.0.3) for high-resolution cellular subclasses annotation based on the reference snRNA-seq dataset^25, 81^.

After predicting the major cell types based on RNA assay, we performed open chromatin peak calling on ATAC assay at pseudo-bulk level with MACS2^82^ using the *CallPeaks* function in Signac. Peaks that overlapped with genomic blacklist regions for the GRCh38 genome were removed, and peak counts were normalized using Latent Semantic Indexing (LSI), including term-frequency inverse-document-frequency (TFIDF). Dimension reductions were performed with singular value decomposition (SVD) of the normalized open chromatin accessibility matrix. The open chromatin accessibility UMAP visualization was created using the second through the 50th LSI components.

To mitigate batch effects, we applied the Harmony (v1.0.0)^83^ algorithm on both snRNA-seq and snATAC-seq data. Then, we created a weighted nearest neighbor (WNN) graph with the *FindMultiModalNeighbors* function of Seurat to represent a weighted combination of both modalities.

### Cell type proportional analysis by scCODA

We applied the single-cell compositional data analysis using scCODA (v0.1.9)^27^, a Bayesian model, to detect the cell type proportional shift across brain regions and conditions. The statistical test and the posterior inclusion probabilities (Pinc) for credible effects were calculated by including either region or sEOAD diagnosis labels as covariates at a false discovery rate (FDR) < 0.05.

### Differential expression analysis

The DEGs between sEOAD and control samples (sEOAD DEGs) were determined for each major cell type across brain regions. VLMC/pericytes and endothelial cells were excluded from the DEG analysis due to low cell counts. First, we conducted DEG analysis using the MAST framework for the genes presented in at least 25% nuclei in either of the two condition groups. Gene expression data underwent log-normalization with a scale factor of 1 ×10^5^. Age, sex, and batch were included in the MAST model as covariates. Second, we implemented a mixed-effect model, utilizing the Libra R package^84^, to account for the random effect of subject origin, which accommodates a zero-inflated negative binomial distribution. The proposed formula of the mixed-effect model is:

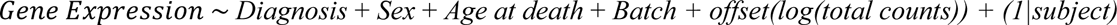

Genes were considered sEOAD DEGs if they met a Bonferroni adjusted p-value threshold of <0.05 in both the MAST and the mixed-effect model, along with an absolute log2FC greater than 0.25 between sEOAD and control in specific cell types.

### Gene Ontology (GO) enrichment analysis

ClusterProfiler (v4.10.0) was used for gene set enrichment analysis to find potential GO BP terms for up- or down-regulated sEOAD DEGs in each cell type within each brain region^85^. The background gene set for this analysis included all genes listed in the GO database. Additionally, we used the *simplify* function to remove redundant enriched GO BP terms based on calculated similarity (similarity score >0.8). The multiple testing correction was performed, and FDR <0.05 was applied.

### Open chromatin peaks annotations

The open chromatin peaks identified in each major cell type were annotated using ChIPseeker^86^ and TxDb.Hsapiens.UCSC.hg38.knownGene using default 3-kb upstream and 3-kb downstream of the TSS. Additionally, we examined the intersection between open chromatin peaks identified in our study and three published datasets: cCRE identified in single nuclei from the adult human brain^4^ and ENCODE Registry of cCRE in the human genome^38^. Brain cell type-specific cis-eQTL data was obtained from Bryois, J. et al.^55^

### Peak-gene linkage analysis

We conducted peak-gene linkage analysis on sEOAD DEGs resulting from the previous step in each cell type using the *LinkPeaks* function in Signac^79^, based on the approach initially described by SHARE-seq^87^. The Spearman correlation coefficient between gene expression and peak accessibility was calculated. Open chromatin peaks within 5 ×10^5^ bp from the gene TSS were included in the model. The GC content, overall accessibility, and peak size were considered in the model as covariates for correcting bias. Benjamini-Hochberg (BH) adjusted p-values were calculated in each region. Only high-confidence positive peak-gene links with BH-adjusted p-value < 0.05 and correlation coefficients > 0.05 were retained for downstream analyses.

### Single-nuclei TF binding motif analysis

We retrieved a comprehensive list of TF binding motif position weight matrices covering 755 TFs from the JASPAR 2020 database (collection = “CORE”, species = “*Homo sapiens*”, access date: March 9th, 2024). The DNA sequence motif information was added to the snATAC-seq assay using the *AddMotifs* function, and the motif activity matrix was calculated using chromVAR implemented in the Signac package^79^.

To prioritize the TFs associated with sEOAD, we identified overrepresented motifs for each set of cCREs linked to sEOAD DEGs in different cell types and brain regions. This involves employing the *FindMotifs* function to conduct a hypergeometric test, evaluating the likelihood of the observed motif frequency occurring by chance relative to a GC content-matched background peak set. The background set was uniquely determined for each set of peaks linked to DEGs in each cell type. Enriched TFs were kept if the TF gene was expressed in over 25% of the corresponding cell type.

Additionally, we identified TFs with differential motif activity by performing differential testing on the chromVAR z-score between sEOAD and controls in each major cell type using the *FindMarker* function at Bonferroni adjusted p-value < 0.05. Footprinting analysis of selected TFs was conducted, accounting for the Tn5 sequencing insertion bias.

### Gene regulatory network inference

We applied the SCENIC+ workflow to construct key TF regulatory networks in specific cell types and conditions^42^. Firstly, the topic modeling of the snATAC-seq data was carried out using pycisTopic to identify the sets of co-accessible regions and the candidate enhancer regions. Next, the motif enrichment analysis on the candidate enhancer regions was carried out using cis-target and DEM algorithms on the cell type- and condition-specific differentially accessible regions. Lastly, the TF regulatory networks were built by calculating the accessible region-gene and TF-gene relationships using a gradient-boosting machine regression method. The retrieved regulatory networks were filtered out if the region-to-gene relationship was negative. The TF-cistrome relationships (Rho) were calculated on the filtered TF regulatory networks, and only the regulons with the top 5% highest positive correlation values and 5% lowest negative correlation values were retained for further downstream analysis.

### Differential cell-cell communication analysis

We employed MultiNicheNet (v1.0.1, last updated on March 20^th^, 2024)^49^, a universal tool to infer the putative context-specific cell-cell communication signals mediated by ligand-receptor pairs across cell type and brain regions in sEOAD and control samples. MultiNicheNet ranks the importance of ligand-receptor pairs based on the ‘pseudo-bulk’ transcriptome profile of each sample. Potential ligands were extracted if expressed in at least 25% of the sender cells within their respective clusters, and the p-value cutoff was set to 0.05. To prioritize highly variable intercellular signals in sEOAD, we highlighted the top 25 differential signals sent and received by selected cell types per experimental group. Furthermore, we applied the *make_ligand_activity_target_plot* function and calculated the regulatory potential scores between prioritized ligands and potential targeted genes within the receiver cell using default parameters.

### Genetic variants selection and LD expansion

Index SNPs were collected from four large-scale LOAD GWA studies^17, 18, 20, 21^. LD expansion was performed using LDlinkR^88^ to identify SNPs with LD R^2^ >0.8 based on the Phase 3 of The 1000 Genomes European ancestry^89^. We performed Fisher’s exact test to assess the enrichment of genetic variants associated with LOAD in cCREs linked to sEOAD DEGs in each cell type across brain regions with a Holm–Bonferroni adjusted p-value <0.05 as the cutoff. The background cCREs set was defined as open chromatin peaks identified in each cell type but not linked to sEOAD DEGs.

### GWAS heritability enrichment

The GWAS heritability enrichment analysis was conducted using stratified LD score regression (sLDSC v1.0.1)^56^. sLDSC calculated the proportion of GWAS heritability attributable to SNPs within a specific genomic feature through a regression model. We obtained summary statistics from four large-scale LOAD GWAS^17, 18, 20, 21^. Additional GWAS summary statistics data were downloaded for 12 immune-mediated and nine psychiatric disorders^57–74^ (**Supplementary Table 3**).

We performed sLDSC analysis in two scenarios. First, the sLDSC analysis was conducted on cCREs linked to sEOAD DEGs in each cell type in each region (Spearman correlation >0.05; BH adjusted p-values <0.05). Second, we applied sLDSC on cell type-specific cCREs. The cell type-specific cCREs were identified using the *FindAllMarkers* function from the Seurat package. This function employed a logistic regression model and adjusted for the library size. We prioritized the peaks exhibiting an Bonferroni adjusted p-value <0.01 and absolute log2FC ≥1 in each cell type as the cell type-specific cCREs.

### Power analysis of differentially expressed genes analysis

To estimate the power of our dataset in detecting alterations in gene expression, we performed a simulation-based power analysis using the R package Hierarchicell^90^. This algorithm takes multiple factors into consideration, including gene dropout rates, intra-individual dispersion, inter-individual variation, variable or fixed number of cells per individual, and the correlation between cells within an individual. We employed *compute_data_summaries* and *power_hierarchicell* functions to simulate and estimate the power with the following parameters: the number of samples in each condition (n = 5, PFC; n = 3, EC; n =3, HIP), the minimal fold change between conditions (rho = 1.5 or 2, two parameters), the p-value cutoff (p = 0.05), mean of the number of cells in each sample of each condition, and number of genes (n = 3000).

### Statistics and reproducibility

All statistical methods and analyses used in this study were described in the Methods, figure legends, or main text as appropriate.

## Data availability

The raw and processed single-nucleus multiome data along with related sample information in this study will be deposited in the Gene Expression Omnibus (GEO) database for public access.

## Code availability

The scripts used to generate the images in this manuscript will be available on GitHub (https://github.com/bsml320/snMultiome_sEOAD). The scripts use openly accessible packages within the R (v.4.3.2) and Python (v3.8.18) environments. The parameters used in each analysis were described in the Results or the Methods section.

## Acknowledgments

Human tissue was obtained from the NIH NeuroBioBank through the University of Miami. We extend our gratitude to the donors and their families for their generous contributions to research. This study was partially supported by grants from the National Institutes of Health (U01AG079847 and R01LM012806). The sequencing data were generated by the UTHealth Cancer Genomics Core funded by the Cancer Prevention and Research Institute of Texas (CPRIT RP180734). A.L. was supported by a training fellowship from the Gulf Coast Consortia on Training in Precision Environmental Health Sciences (TPEHS) Training Grant (T32ES027801). The funders had no role in the study design, data collection, analysis, decision to publish, or preparation of the manuscript.

## Contributions

Conceptualization: Z.Z.; methodology: A.L., X.C., L.M.S., and Z.Z.; formal analysis: A.L., C.C., N.E., and A.M.M.; tissue and data generation: X.C., T.S., M.Y., P.S., and C.S.; writing—original draft: A.L., C.C., A.M.M., N.E., T.S., D.G., B.S.F., and X.C.; writing—review and editing: Z.Z., L.M.S., B.S.F. and C.S.; visualization: A.L., C.C., and N.E.; funding acquisition: Z.Z.; supervision: Z.Z., L.M.S. All authors read and approved the final manuscript.

## Corresponding author

Correspondence to Zhongming Zhao

---

**Extended Data Fig. 1:**
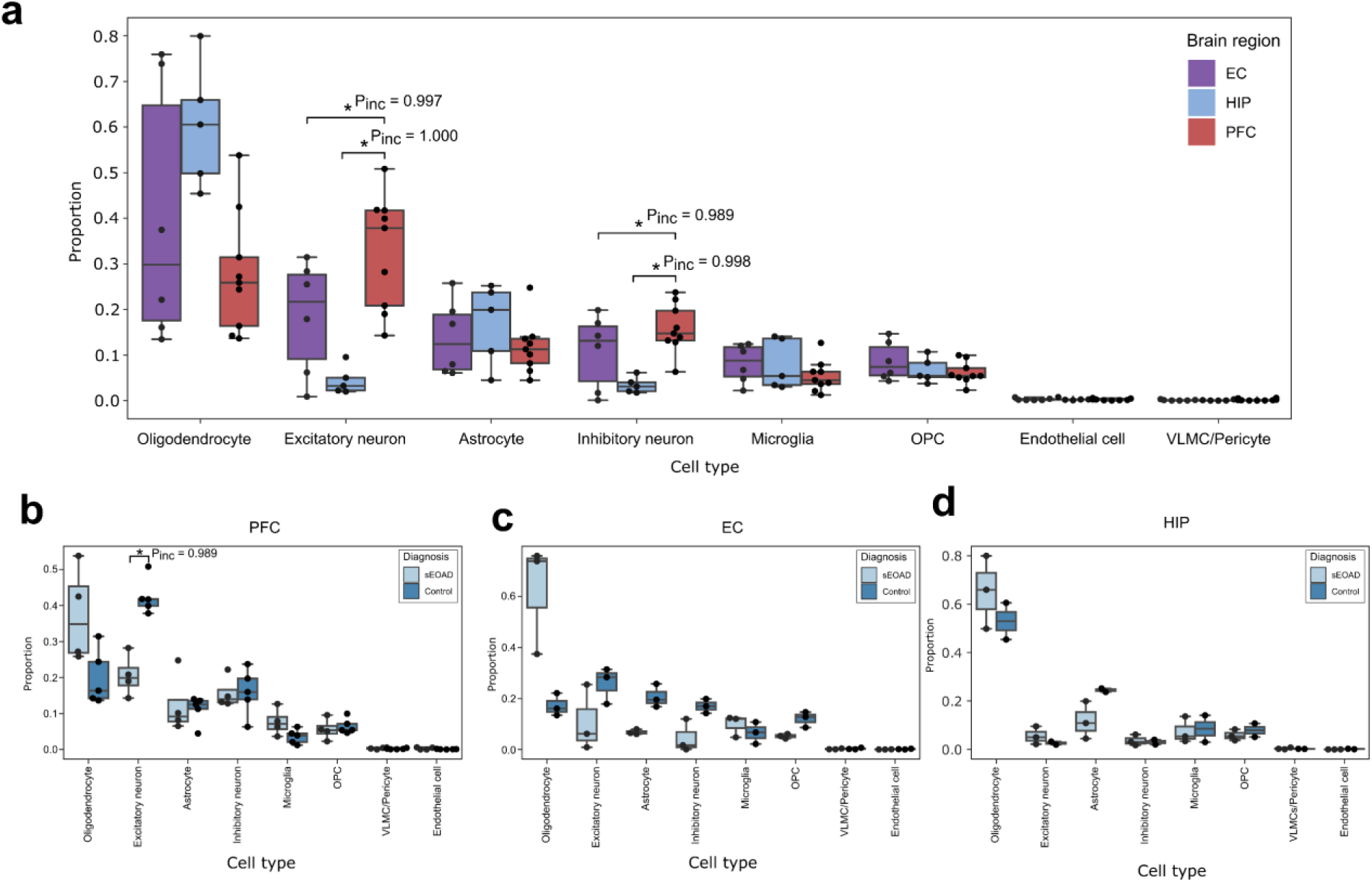
Differential cell type composition across brain regions and conditions. **a,** Major cell type compositional changes comparing samples from the entorhinal cortex (EC), hippocampus (HIP), and prefrontal cortex (PFC). **b-d,** Major cell type compositional changes between sEOAD and control samples in (**b**) PFC, (**c**) EC, and (**d**) HIP. Box boundaries and lines in box plots correspond to the interquartile range (IQR) and median, respectively. Statistically credible changes in cell types, as tested with scCODA, are marked with *Pinc (posterior inclusion probability), indicating the significance of these changes.

**Extended Data Fig. 2:**
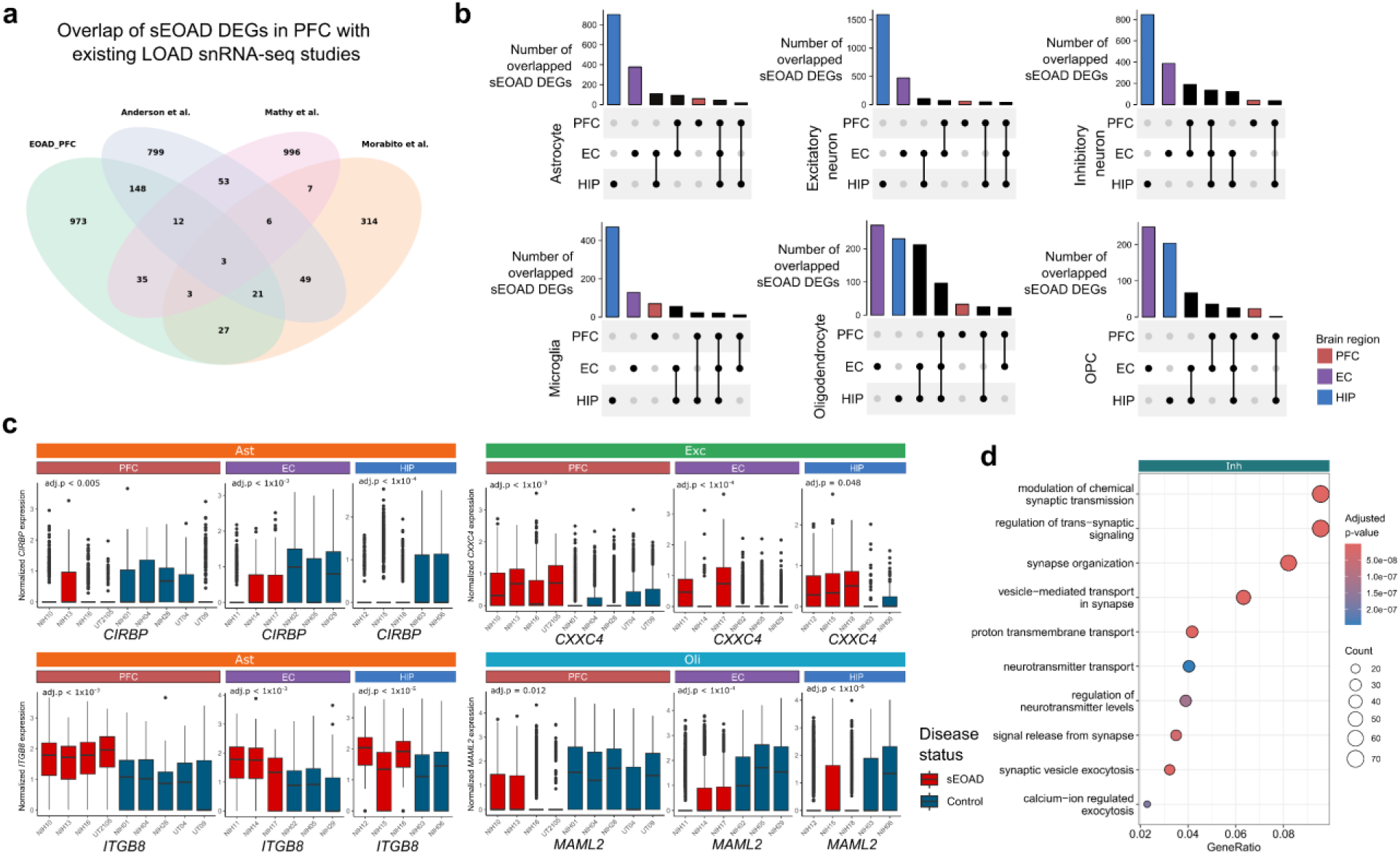
Overlap of cell type-specific sEOAD DEGs across brain regions and studies. **a,** Venn diagram showing the overlap of sEOAD DEGs in the prefrontal cortex (PFC) identified in current study with signals reported by existing late-onset Alzheimer’s Disease (LOAD) single-nuclei RNA-sequencing studies. Three DEGs were consistent across all studies, including *MRAS* in astrocytes and *CNORDC1* and *TP53TG3* in oligodendrocytes. **b,** The number of overlapped sEOAD differential expression genes (sEOAD DEGs) in each cell type across brain regions with colors indicating brain regions. **c,** sEOAD DEGs consistently identified in all three brain regions associated with specific pathways: circadian clock regulation (top left, *CIRBP* in astrocyte), extracellular matrix organization (top right, *ITGB8* in astrocyte), Wnt-signaling pathway (bottom left, *CXXC4* in excitatory neurons), NOTCH signaling pathway (bottom right, *MAML2* in oligodendrocytes). The x-axis indicates sample identification, and the color of the box indicates the diagnosis label. Box boundaries and lines correspond to the interquartile range (IQR) and median, respectively. **d,** The top 10 enriched Gene Ontology biological processes (GO BP) terms for consistently downregulated sEOAD DEGs in inhibitory neurons (number of genes = 110) across brain regions.

**Extended Data Fig. 3:**
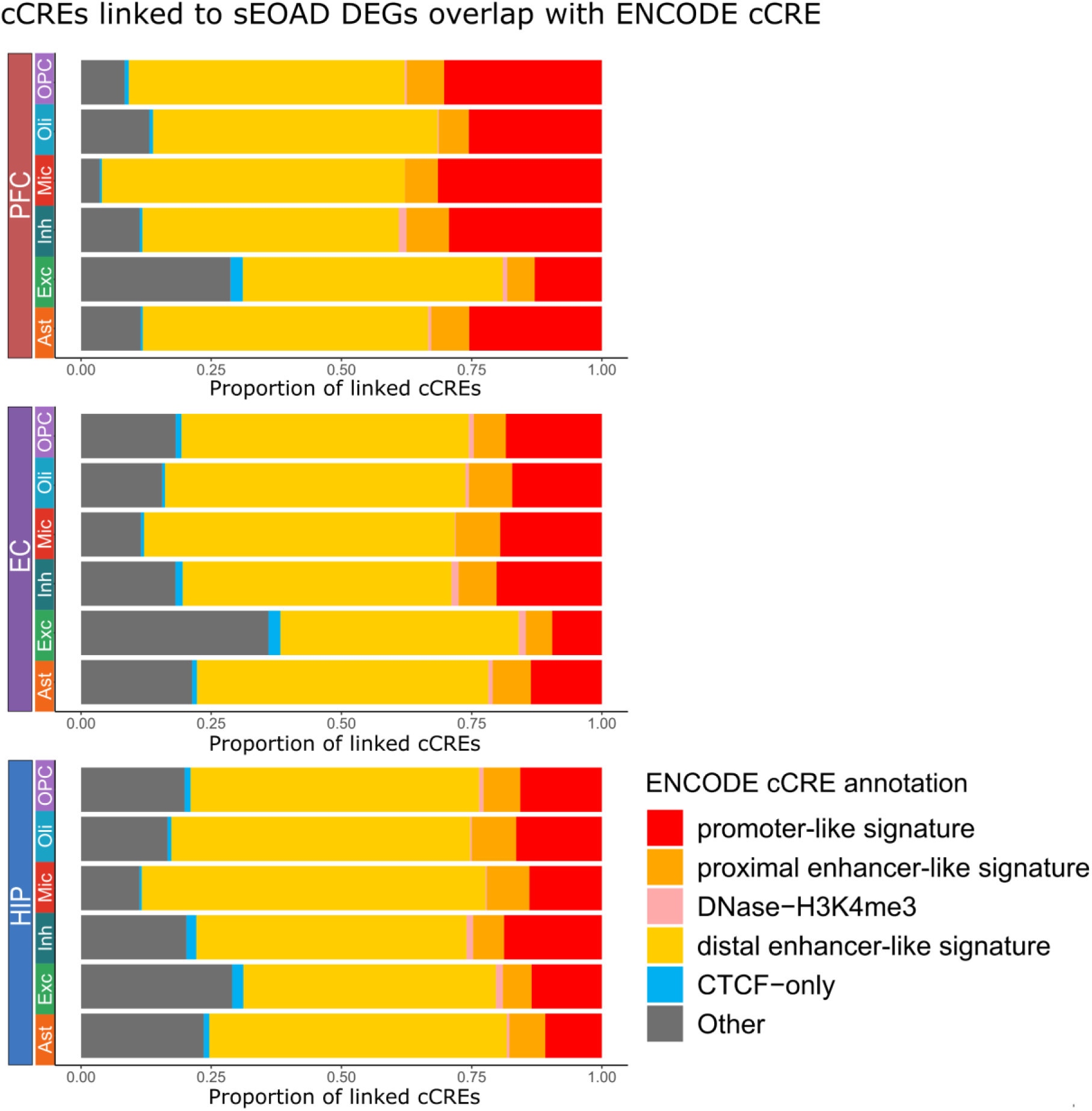
ENCODE annotation of *cis-*regulatory elements linked to sEOAD DEGs by cell type and brain region. Stacked bar plots displaying the percentage of candidate cis-regulatory elements linked to sEOAD differential expression genes (DEGs) in each cell type and region that overlapped with ENCODE cCREs annotated promoters and enhancers.

**Extended Data Fig. 4:**
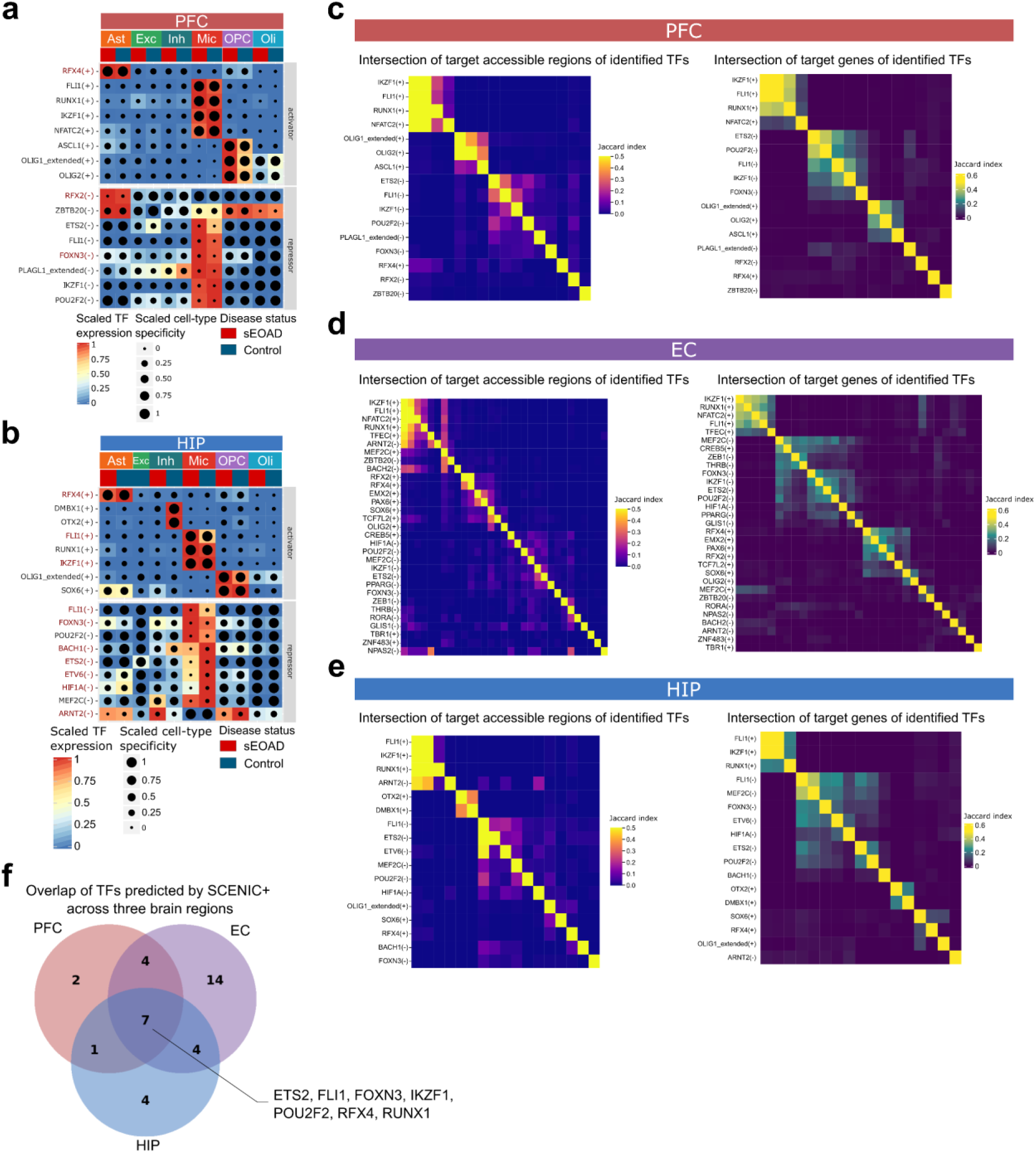
Cell type-specific transcription factor regulatory circuities in the prefrontal cortex and hippocampus of sEOAD. **a,b,** Heatmap/dot plot showing SCENIC+ analysis results for (**a**) the prefrontal cortex (PFC) or (**b**) the hippocampus (HIP), categorized by cell type and diagnosis label (x-axis). The gene expression of identified transcription factors (TFs) is presented on a color scale, while the cell type-specificity of the regulon is presented by dot size. **c-e**, Jaccard heat maps showing the intersection of target regions (left) and target genes (right) of the identified TFs in (**c**) PFC, (**d**) EC, and (**e**) HIP. **f,** Venn diagram depicting the number of overlaps of transcription factors in the PFC, EC, and HIP.

**Extended Data Fig. 5:**
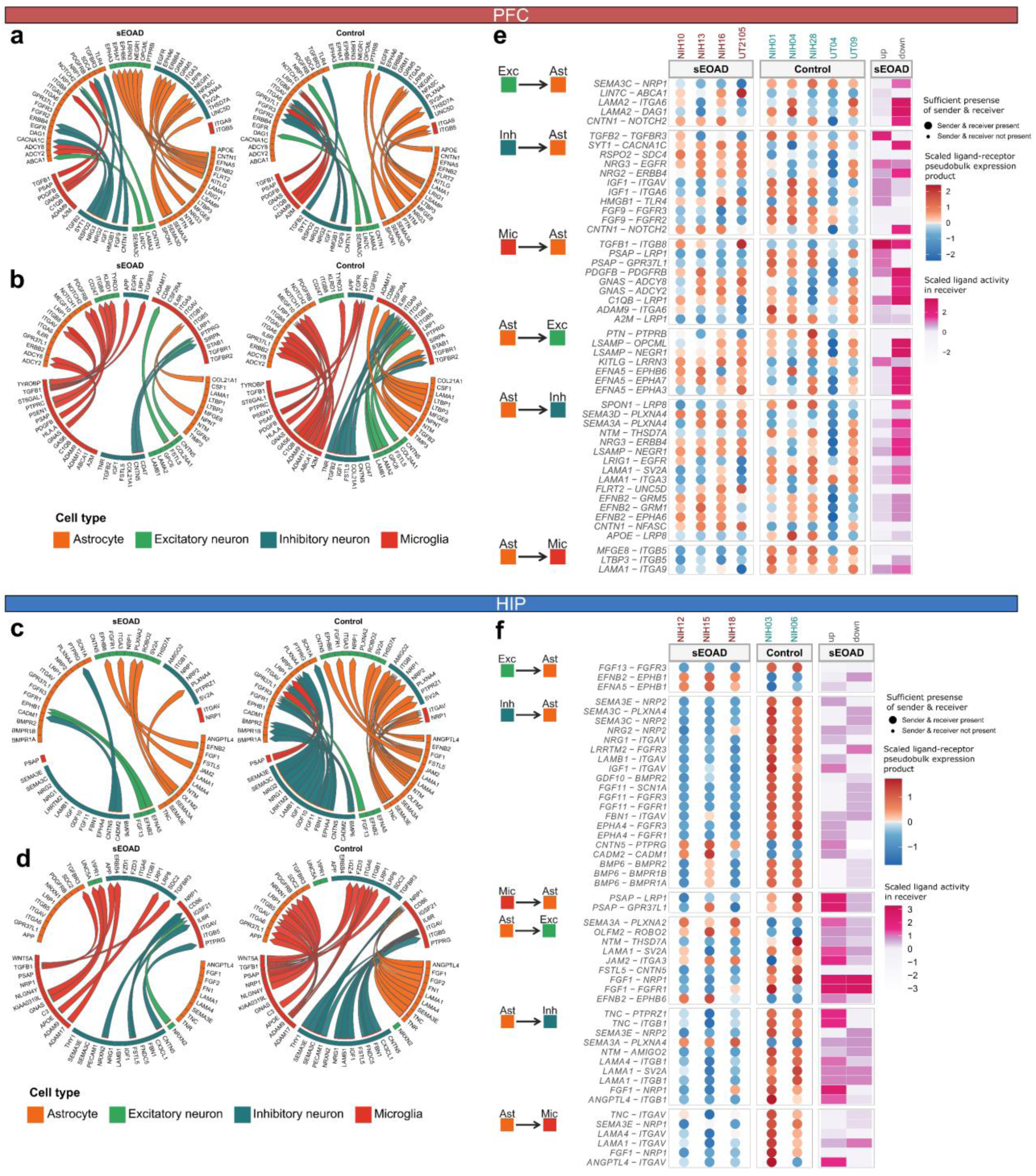
Comparative analysis of dysregulated cell-cell communication in PFC and HIP between sEOAD and controls. **a,** Top 25 differential ligand-receptor signals received and sent by astrocytes in the prefrontal cortex (PFC) between sEOAD and control groups. **b,** Top 25 differential ligand-receptor signals received and sent by microglia in the PFC between sEOAD and control groups. **c,** Top 25 differential ligand-receptor signals received and sent by astrocytes in the hippocampus (HIP) between sEOAD and control groups. **d,** Top 25 differential ligand-receptor signals received and sent by microglia in the HIP between sEOAD and control groups. The arrowhead represents the direction of the signal from the sender cell type to the receiver cell type. **e-f**, The heatmap depicting the regulatory potential scores (purple) of selected dysregulated ligand-receptor pairs in receiver cell types within (**e**) PFC or (**f**) HIP. The dot plots show the scaled ligand-receptor pseudobulk expression in each sample, with dot size indicating whether a sample had at least ten cells for astrocytes.

**Extended Data Fig. 6:**
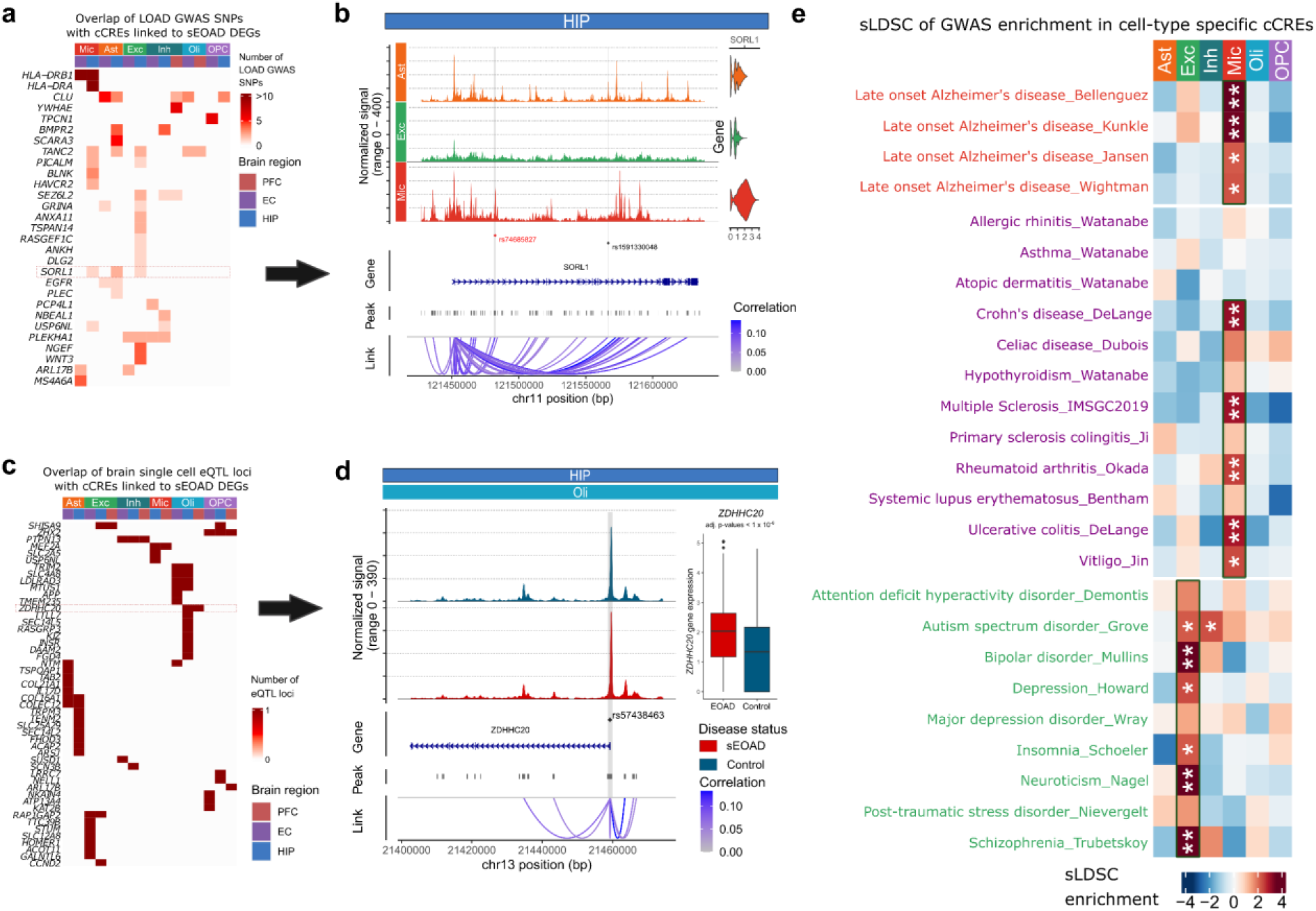
GWAS enrichment in cell type specific cCREs and variants prioritization. **a,** Heatmap displaying the sEOAD differentially expressed genes (DEGs) with associated candidate *cis-*regulatory elements(cCREs) containing LOAD GWAS loci. **b,** SNP rs74685827, as reported in Bellenguez *et al.,* overlaps with a microglia-specific cCRE linked to dysregulated *SORL1* gene expression in hippocampus (HIP). Its associated SNP, rs1591330048, overlaps with a cCRE linked to dysregulated *SORL1* gene expression in astrocytes in HIP. The cCREs linked to sEOAD DEG contain LOAD GWAS loci and are highlighted in gray. **c,** Heatmap shows the cCRE linked to sEOAD DEGs containing single-cell expression quantitative trait loci (eQTL). **d,** The promoter region of *ZDHHC20* in oligodendrocytes contains the single-cell eQTL SNP rs57438463. The box plot depicts that the *ZDHHC20* gene was significantly upregulated in oligodendrocytes in HIP samples of sEOAD patients. **e,** Heatmap showing the enrichment of risk variants associated with LOAD, immune-mediated traits, and psychiatry disorders from GWAS in cell type specific peaks (*FDR < 0.05, **FDR < 0.01).

